# Assembly of Long Error-Prone Reads Using Repeat Graphs

**DOI:** 10.1101/247148

**Authors:** Mikhail Kolmogorov, Jeffrey Yuan, Yu Lin, Pavel. A. Pevzner

## Abstract

The problem of genome assembly is ultimately linked to the problem of the characterization of all repeat families in a genome as a *repeat graph*. The key reason the *de Bruijn graph* emerged as a popular short read assembly approach is because it offered an elegant representation of all repeats in a genome that reveals their mosaic structure. However, most algorithms for assembling long error-prone reads use an alternative *overlap-layout-consensus (OLC)* approach that does not provide a repeat characterization. We present the Flye algorithm for constructing the A-Bruijn (assembly) graph from long error-prone reads, that, in contrast to the *k*-mer-based de Bruijn graph, assembles genomes using an alignment-based A-Bruijn graph. In difference from existing assemblers, Flye does not attempt to construct accurate contigs (at least at the initial assembly stage) but instead simply generates arbitrary paths in the (unknown) assembly graph and further constructs an assembly graph from these paths. Counter-intuitively, this fast but seemingly reckless approach results in the same graph as the assembly graph constructed from accurate contigs. Flye constructs (overlapping) contigs with possible assembly errors at the initial stage, combines them into an accurate assembly graph, resolves repeats in the assembly graph using small variations between various repeat instances that were left unresolved during the initial assembly stage, constructs a new, less tangled assembly graph based on resolved repeats, and finally outputs accurate contigs as paths in this graph. We benchmark Flye against several state-of-the-art Single Molecule Sequencing assemblers and demonstrate that it generates better or comparable assemblies for all analyzed datasets.

## INTRODUCTION

The problem of genome assembly is ultimately linked to the *repeat characterization problem*, the compact representation of all repeat families in a genome as a *repeat graph* (Pevzner et al., 2004). Long read technologies have not made the repeat characterization problem irrelevant. Instead, they have simply shifted the focus from short repeats to longer repeats comparable in length to the median SMS read size; e.g., Kamath et al., 2017 analyzed many bacterial genomes that existing SMS assemblers failed to assemble into a single contig. Since even bacterial (let alone, eukaryotic) genomes have long repeats, SMS assemblers currently face the same challenge that short read assemblers faced a decade ago, albeit at a different scale of repeat lengths. To address this assembly challenge, SMS technologies are often complemented by contact (Ghurye et al., 2017) and optical (Weissensteiner et al., 2017) mapping data.

The *de Bruijn graph* emerged as a popular short read assembly approach because it offered an elegant representation of all repeats in a genome. However, most long read assemblers use an alternative *overlap-layout-consensus (OLC)* approach that requires an all-against-all comparison of reads, does not yield a natural representation of all repeats in a genome, and does not provide optimal repeat resolution (Koren et al., 2012, Chin et al., 2013, Berlin et al., 2015, Chin et al., 2016, Li, 2016, Koren et al., 2017). Recent studies of SMS assemblies attempted to integrate the OLC approach by constructing the alternative de Bruijn graph of reads (referred to as the *assembly graph*). Although these approaches improved on the OLC-only bacterial assemblies (Lin et al., 2016, Kamath et al., 2017), the de Bruijn graph constructed from error-prone SMS reads is very noisy making it difficult to collapse various instances of the same repeat into a single path in the assembly graph. Developers of OLC approaches realized the importance of repeat characterization and attempted to improve the *best overlap graph (BOG)* representation (Miller et al., 2008) by developing the OLC-based *Bogart graph* for representing repeats (Koren et al., 2017). However, the Bogart graph is quite different from the repeat graph and, as acknowledged by the developers of the Canu assembler (Koren et al., 2017), it is still error-prone (see Kamath et al., 2017 for a discussion on the differences between the de Bruijn-based and the OLC-based repeat representation). As an alternative to these overlap graphs, which are riddled with unnecessary edges, the *string graph* (Meyers, 2005) approach (implemented in the Miniasm assembler) attempts to represent unique segments of a genome as non-branching paths. However, as discussed in Kamath et al., 2017, the string graph does not achieve an optimal repeat resolution either.

*The challenge of constructing the assembly graph from long reads.* Most short read assemblers construct the de Bruijn graph based on all *k*-mers in reads and further transform it into an *assembly graph* using various *graph simplification* procedures. This approach collapses multiple instances of the same repeat into a single path in the assembly graph and represents a genome as a *genome tour* that visits each edge in the assembly graph. However, in the case of SMS reads, the key assumption of the de Bruijn graph approach (that most *k*-mers from the genome are preserved in multiple reads) does not hold even for short *k*-mers, let alone for long *k*-mers (e.g., *k* = 1000). As a result, various issues that have been addressed in short read assembly (e.g., how to deal with the fragmented de Bruijn graph, how to transform it into an assembly graph, etc.) remain largely unaddressed in the case of the de Bruijn graph approach to SMS assemblies

Aggressive approaches to repeat resolution often produce misassemblies while conservative approaches lead to fragmented assemblies (Kamath et al., 2017). Construction of an assembly graph is an important step towards finding the balance between aggressive and conservative approaches since it allows one to distinguish repeats that can be resolved from those that cannot. Constructing an assembly graph for SMS reads improves repeat resolution as compared to various OLC assemblers (Kamath et al., 2017) because it is better suited for finding this balance. Also, since assembly graphs represent a special case of *breakpoint graphs* used for genome rearrangement studies (Lin et al., 2014), they are well suited for the analysis of genome variations using SMS reads (Chaisson et al., 2015, Nattestad et al., 2017).

Here, we describe the Flye algorithm for constructing assembly graphs from polished contigs rather than from error-prone reads (as in HINGE), thus making the graph more accurate (see Appendix “Short read assemblers and the challenge of repeat resolution” for a discussion on contig assembly in the case of short read sequencing). Flye also complements HINGE by introducing a new algorithm that uses small differences between repeat copies to resolve *unbridged* repeats that are not spanned by any reads.

Flye is built on top of the ABruijn assembler (Lin et al., 2016), which generates *accurate* overlapping contigs but does not reveal the repeat structure of the genome. In contrast to ABruijn, Flye initially generates inaccurate overlapping contigs (i.e., contigs with potential assembly errors) and combines these *initial* contigs into an accurate assembly graph that encodes all possible assemblies consistent with the reads. Flye further resolves unbridged repeats in the assembly graph thus constructing a new, less tangled assembly graph, and finally outputs accurate *final* contigs formed by paths in this graph. We benchmarked Flye against several state-of-the-art SMS assemblers using various datasets and demonstrated that it generates accurate assemblies while also providing insights into how to plan additional experiments (e.g., using contact or optical maps) to finish the assembly.

## METHODS

### Repeat graphs

Repeats in a genome are often represented as pairwise local alignments between various regions of the genome and visualized as two-dimensional *dot plots* representing all alignment paths. This pairwise representation is limited since it does not contribute to solving the repeat characterization problem. In contrast, the *repeat graph* (Pevzner et al., 2004) compactly represents all repeats in a genome and reveals their mosaic structure. The construction of an assembly graph represents a special case of the repeat graph construction problem, which we discuss below.

Consider a tour *T*=*v*_1_, *v*_2_,…*v*_n_ of length *n* visiting all vertices of a directed graph *G*. We say that the *i*-th and *j*-th vertices in the tour *T* are *equivalent* if they correspond to the same vertex of the graph (i.e., *v_i_*=*v_j_*). The set of all pairs of equivalent vertices forms a set of points (*i,j*) in a two-dimensional grid that we refer to as the *repeat plot Plot_T_(G)* of the tour *T* (Figure 1). The transformation of a tour *T* traversing a *known* graph *G* into the repeat plot *Plot_T_(G)* is a simple procedure. Below, we address a reverse problem that is at the heart of genome assembly, repeat characterization, and synteny block construction: given an arbitrary set of points *Plot,* in a two-dimensional grid, find a graph *G*=*G(Plot)* and a tour *T* in this graph such that *Plot* = *PlotT(G).* In the case of repeat characterization, *Plot* consists of points traversed by paths representing all local self-alignments (up to a certain stringency) of a genome against itself. Each self-alignment reveals two instances of a repeat corresponding to genomic segments *x* and *y* (*x* and *y* are called the *spans* of the alignment). Given a genome of length *n* and a set of its pairwise self-alignments *Plot*, the goal is to construct a graph *G* and a tour *T* of length *n* in this graph (each segment of the genome corresponds to a subpath of the graph traversed by the tour) such that *Plot* = *Plot_T_(G)* and for each alignment with spans *x* and *y,* paths in the graph corresponding to segments *x* and *y* coincide.

### Generating all local self-alignments of a genome

Our goal is to represent all long repeats in a genome as a repeat graph (Pevzner et al., 2004). Constructing the de Bruijn graph based on *long k*-mers will not solve this problem since the differences between imperfect repeat copies mask the repeat structure of the genome. Constructing the de Bruijn graph based on *short k*-mers will not solve this problem due to the presence of repeating short *k*-mers within long repeats (these *k*-mers lead to a tangled repeat graph). Thus, while all long similar segments can be generated based on pairwise alignments and combined into a repeat plot *Plot*, it is unclear how to solve the reverse problem of combining them into the repeat graph *G(Plot)* of the genome. Although Pevzner et al., 2004 described various graph simplification procedures (that are now at the heart of various short read assemblers) to address this problem for short reads, it is not clear how to generalize these procedures to make them applicable to SMS assemblies. Below we describe how the concept of a punctilious repeat graph (introduced in Pevzner et al., 2004) can be modified for assembling SMS reads.

### From local alignments to a punctilious repeat graph

Let *Alignments*=*Alignments*_k_(*Genome*) be the set of all pairwise local alignments of length at least *k* between various segments of a genome *Genome*. Given a set of self-alignments *Alignments* of a genome *Genome*, we construct the *punctilious repeat graph* G*(Genome, Alignments*) by representing *Genome* as a path consisting of |*Genome*| vertices (Figure 1) and simply “gluing” each pair of vertices (positions in the genome) that are aligned against each other in one of the alignments in *Alignments* (Pevzner et al., 2004). Gluing vertices *v* and *w* amounts to substituting them by a single vertex that is connected by edges to all vertices that either vertex *v* or vertex *w* is connected to. We consider *branching vertices* (i.e., vertices with either in-degree or out-degree differing from 1) in the resulting graph and substitute each *non-branching path* of length *m* between them by a single edge of length *m*. Edges in the punctilious assembly graph are classified into *long* (longer than a predefined threshold *d*) and *short* (Figure 1).

**Figure 1.**
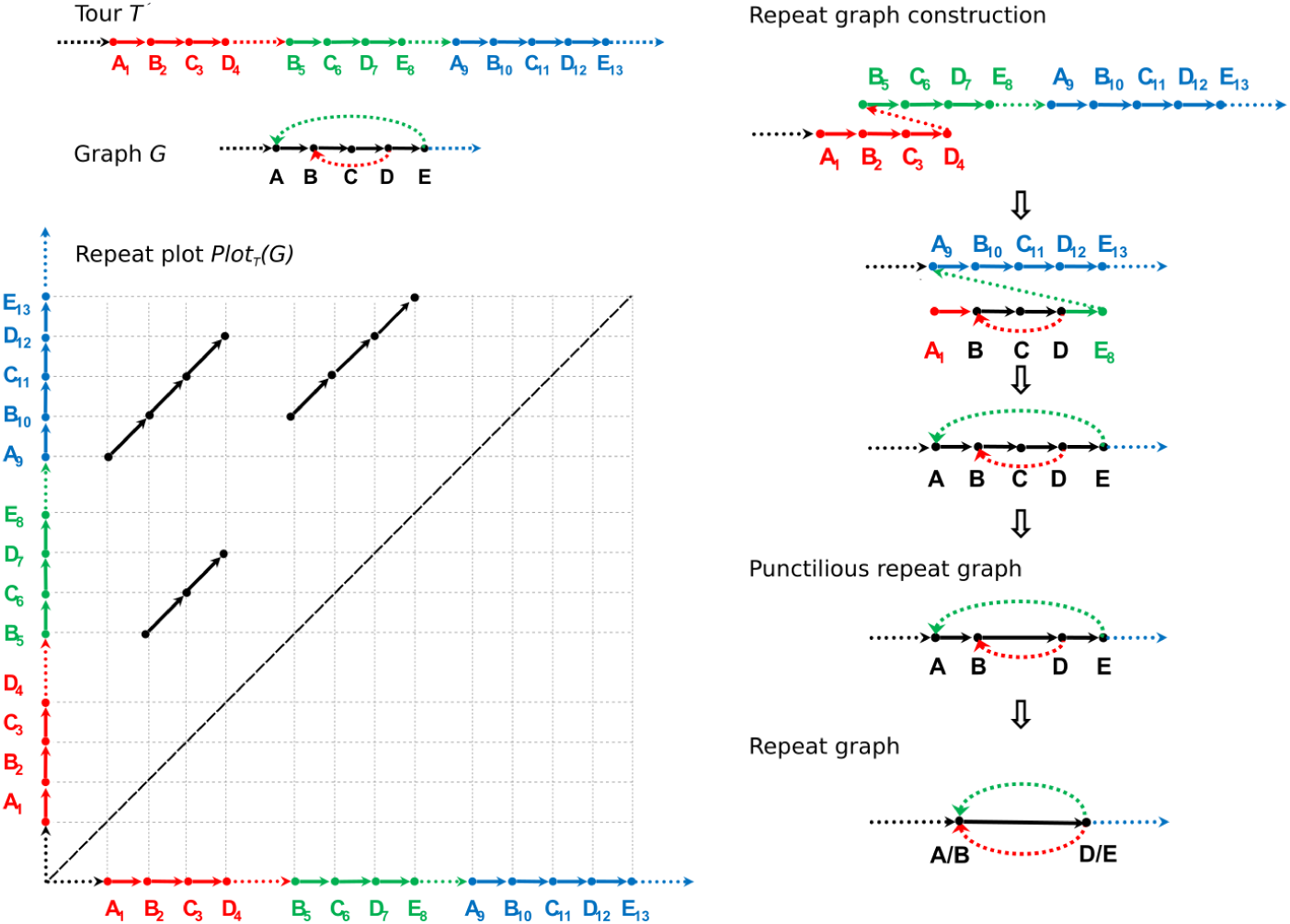
Constructing the repeat plot of a tour in a graph (left) and constructing the repeat graph from a repeat plot (right) (Left) A tour 
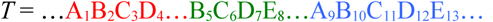
 in a graph *G* with red, green, and blue instances of a repeat. Dots represent multiple vertices that appear before, between, and after these three instances of the repeat. The repeat plot *PlotT(G)* consists of three diagonals representing the three instances of the repeat in the tour. The trivial self-alignment of the entire genome against itself is shown by the dotted diagonal (since the repeat plot is symmetric, the points below this diagonal are not shown). Since vertex A in the graph is visited twice in the tour *T*, it results in a single point (1,9) in *Plot_T_(G)*. Vertex B results in points (2,5), (2,10), and (5,10), vertex C results in points (3,6), (3,11), and (6,11), vertex D results in points (4,7), (4,12), and (7,12), and vertex E results in the point (8,13). (Right) Constructing the punctilious repeat graph from the repeat plot by gluing vertices with indices *i* and *j* for each point (*i,j*) in the repeat plot. Each non-branching path in the graph is substituted by a single edge with length equal to the number of edges in this path. The lengths of the short edges (A,B) and (D,E) in the resulting graph are equal to 1 and the length of the long edge (B,D) is equal to 2 (for the edge length threshold *d*=1). The punctilious repeat graph (second graph from the bottom) is transformed into the repeat graph (bottom-most graph) by contracting short edges (A,B) and (D,E).

Figure 1 presents a simple repeat plot where the ending positions of various pairwise alignments are coordinated. In reality, they are typically not coordinated, e.g., three alignment paths corresponding to three repeat copies starting at position *x, y,* and *z* hardly ever start at points (*x,y*), (*x,z*), and (*y,z*) in the repeat plot. Because each repeat with *m* copies results in 
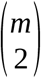
 local alignments, there will be 
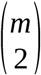
 gluing operations for the starting positions of this repeat and the same number of gluing operations for the ending positions of this repeat. Note that each of these operations may form a new branching vertex in the punctilious repeat graph. For example, gluing the endpoints of the three diagonals in Figure 1 results in the branching vertices A, B, C, and D in the graph. Punctilious repeat graphs of real genomes often contain many branching vertices making it difficult to compactly represent repeats. We address this challenge by transforming the punctilious repeat graph into a simpler repeat graph.

### From punctilious repeat graph to repeat graph

Since endpoints of alignment paths representing the same repeat are not coordinated among all pairwise alignments of this repeat, these uncoordinated alignments result in a complex repeat graph with many short edges (shorter than a predefined length threshold *d*). The *repeat graph G*(*Genome, Alignments, d*) is defined as the result of *contracting* all short edges in the punctilious repeat graph (Figure 1). The contraction of an edge is simply the gluing the endpoints of this edge, followed by the removal of the loop-edge resulting from this gluing. Since the genome represents a tour visiting all edges in the repeat graph, we define the multiplicity of an edge in the repeat graph as the number of times this edge is traversed in the tour. Edges of multiplicity 1 are called *unique edges* and edges of multiplicity more than 1 are called *repeats*.

### An alternative approach to constructing repeat graphs

The described approach, although simple in theory, results in various complications in the case of real genomes (see Pevzner et al., 2004), particularly in the case of inconsistent pairwise alignments (see Appendix “Consistent alignments”). In difference from the approach in Pevzner et al., 2004, Flye constructs an *approximate* version of the repeat graph that focuses on the endpoints of alignments rather than the entire alignment paths, resulting in simpler repeat graphs that are better suited for assembling SMS reads.

Some branching vertices in the repeat graph have arisen from the contraction of multiple vertices in the punctilious repeat graph into a single branching vertex in the repeat graph, e.g., vertices A and B were contracted into a single vertex A/B in the repeat graph in Figure 1. Consider the set of all vertices in the punctilious repeat graph that gave rise to branching vertices in the repeat graph (vertices A, B, D, and E in Figure 1) and let *Endpoints* = *Endpoints*(*Genome, Alignments, d*) be the set of all positions in the genome that gave rise to these vertices (*Endpoints*={1,2,4,5,7,8,9,10,12,13} in Figure 1). This set of vertices forms a set of short contiguous genomic segments that contain the endpoints of all pairwise alignments in *Alignments.* As an example in Figure 1, the set *Endpoints* forms segments [1,2], [4,5], [7,8,9,10], and [12,13].

**Figure 2.**
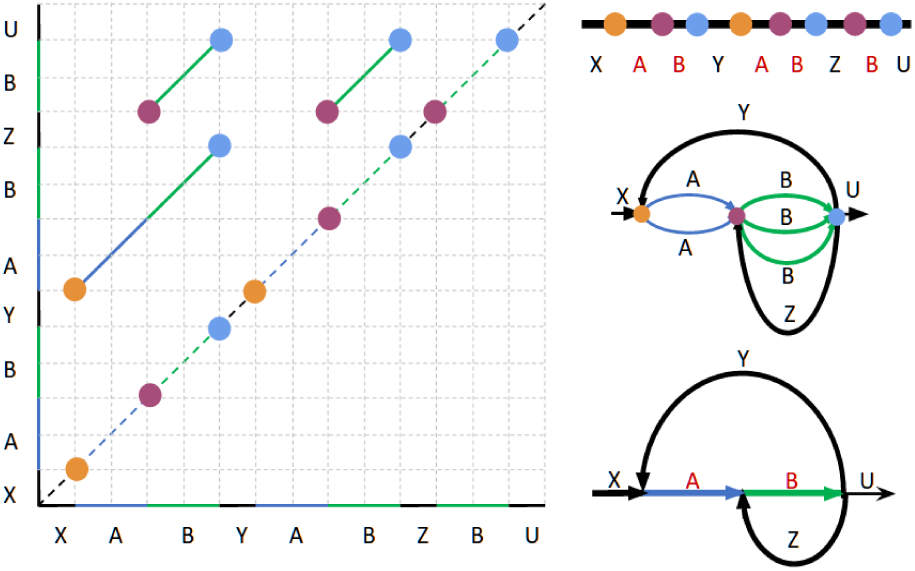
Repeat graph construction from local alignments within a genome. (Left) Alignment paths for all local self-alignments within a genome XABYABZBU (with three instances of a mosaic repeat: AB, AB, and B) represented as a repeat plot. The self-alignment of the entire genome is shown by the dotted diagonal. Alignment endpoints are clustered together if their projections on the main diagonal coincide or are close to each other (clusters of closely located endpoints for *d*=0 are painted with the same color). For example, rightmost endpoints (shown in blue) of all three alignments form a single cluster because every two of them share the same projection on the main diagonal. This clustering reveals three clusters (yellow, purple, and blue) with eight endpoints (Top right) Projections of the clustered endpoints on the main diagonal define eight vertices of the repeat graph. (Middle right). Gluing endpoints that belong to the same clusters. (Bottom right) Gluing parallel edges in the resulting graph (parallel edges are glued if there exists an alignment between their sequences), which results in the approximate repeat graph.

Flye approximates the set *Endpoints* by recruiting all horizontal and vertical projections of endpoints of alignments from *Alignments* to the main diagonal in the repeat plot. Figure 2 presents three alignments resulting in eight projected points on the main diagonal. Two alignment endpoints are *close* if their projections on the main diagonal are located within distance threshold *d* (including the case when a vertical projection of one endpoint coincides or is close to a horizontal projection of another endpoint). Flye clusters close endpoints together based on single linkage clustering, resulting in three clusters with eight endpoints in Figure 2 (clusters for *d*=0 are painted with the same color). Flye further approximates the repeat graph by gluing endpoints that belong to the same clusters as shown in Figure 2. However, the repeat graph constructed based on the clustering of closely located endpoints may differ from the repeat graph constructed based on the punctilious assembly graph. Appendix “Inconsistent alignments result in incorrect clustering-based repeat graph” illustrates that mosaic repeat structures and inconsistencies of local alignments may result in an “incorrect” clustering-based repeat graph. Below we explain how Flye iteratively extends the set *Endpoints* to address this complication.

### Extending the set of alignment endpoints

Given a set of alignments *Alignments*, Flye first projects their endpoints onto the main diagonals in the repeat plot, resulting in the set *Endpoints*. Each point in an alignment-path in the |*Genome*|×|*Genome*| grid has two projections (horizontal and vertical) on the main diagonal. Note that projections of some internal points in an alignment path may belong to *Endpoints*; for example, both projections of the middle-point of the longest alignment-path in Figure 2 belong to *Endpoints*. Such internal points should be re-classified as new alignment endpoints (essentially by breaking an alignment into two parts) to avoid inconsistencies during the construction of the repeat graph. Below we explain how to extend the set *Endpoints* to address this complication.

A point in an alignment-path is called a *valid point* if both its projections belong to *Endpoints,* and an *invalid point* if only one of its projections belongs to *Endpoints* (see Appendix: “Inconsistent alignments result in incorrect clustering-based repeat graph” for an example of an invalid point). Invalid points complicate the task of generating a new set of alignment endpoints (by sub-partitioning the alignment paths) since alignment endpoints have two rather than a single projection on the main diagonal. Flye thus iteratively adds the missing projection for each invalid point to the set *Endpoints* on the main diagonal until there are no invalid points left. Afterwards, it clusters points in *Endpoints* that are less than *d* nucleotides apart using single linkage clustering. Each resulting cluster corresponds to a segment between its minimal and maximal positions on the main diagonal and these segments form a set *EndpointSegments*. Two segments from *EndpointSegments* are *equivalent* if there exists a valid point in one of the alignments such that one of its projections to the main diagonal falls into the first segment and another falls into the second segment.

Flye selects a middle point from each segment in *EndpointSegments* and glues the two middle points for every pair of equivalent segments. Finally, it glues together parallel edges (edges that start and end at the same vertices) if the genome segments corresponding to these edges are aligned in *Alignments*, i.e., if there exists an alignment with its *x*- and *y*-spans overlapping both these segments.

### From the repeat graph of a genome to the assembly graph of contigs

ABruijn constructs a set of contigs *Contigs* by implicitly “extending” paths in an (unknown) assembly graph. When the path extension process reaches a branching vertex, ABruijn decides whether to continue or to stop the path extension (as described in Lin et al., 2016) in order to avoid assembly errors. Since ABruijn does not know the exact locations of branching vertices, it extends the path beyond the branching vertex by at least a predefined threshold of *minOverlap* nucleotides even in the case that it decides to stop the path extension process. As a result, various contigs constructed by ABruijn overlap. Flye uses these overlaps for assembly graph construction and concatenates all contigs in *Contigs* (separated by delimiters) in an arbitrary order into a single sequence *Contigs*.* It further constructs the *assembly graph* as the repeat graph *Graph*(*Contigs*, Alignments, d*) of the concatenate. Pevzner et al., 2004 demonstrated that the assembly graph of reads approximates the repeat graph of the genome. Thus, the assembly graph constructed from *accurate* contigs (which can be viewed as virtual reads) also approximates the repeat graph of the genome. Below we discuss the construction of assembly graphs from inaccurate contigs.

### Constructing an accurate assembly graph from misassembled contigs

In contrast to all existing SMS assemblers that invest significant effort into making sure that generated contigs are correctly assembled (that they represent subpaths of the genomic tour in the assembly graph), Flye does not attempt to construct accurate contigs (Flye contigs instead represent arbitrary paths in the assembly graph). However, it constructs an accurate assembly graph from inaccurate contigs.

Flye randomly walks in the (unknown) assembly graph to generate random paths from this graph. Each non-chimeric read from *Reads* defines a subpath of a genomic tour in an (unknown) assembly graph. Flye extends this path by switching from the current read to *any* other overlapping read rather than a *carefully chosen* overlapping read like in Lin et al., 2016. Thus, the contig extension is greatly speed-up since it merely selects a read that shares a sufficiently long common *jump*-subpath with the current read (Lin et al., 2016) and avoids a time-consuming test to check if this selection is correct.

Since the resulting FlyeWalk algorithm (Figure 3) does not invoke the contig correctness check, it constructs paths (sequence of overlapping reads) that do not necessarily follow the genome tour through the assembly graph. Although it may appear counter-intuitive that inaccurate contigs constructed by FlyeWalk result in an accurate assembly graph, note that inaccurate contigs in the de Bruijn graph (a special case of the assembly graph) certainly result in an accurate assembly graph. Indeed, an assembly graph constructed from arbitrary paths in a de Bruijn graph is the same as the original de Bruijn graph (as long as these paths include all *k*-mers from the assembly graph). See Appendix “Flye constructs an accurate assembly graph from inaccurate FlyeWalk contigs.”

Figure 4, left, presents a rather complex assembly graph of the SMS reads from an *E. coli* strain NCTC9964 (EC9964 dataset described in the Appendix “Information about the BACTERIA datasets”). Flye further untangles this graph into a graph with just six edges (Figure 4, middle) as described below.

**Figure 3.**
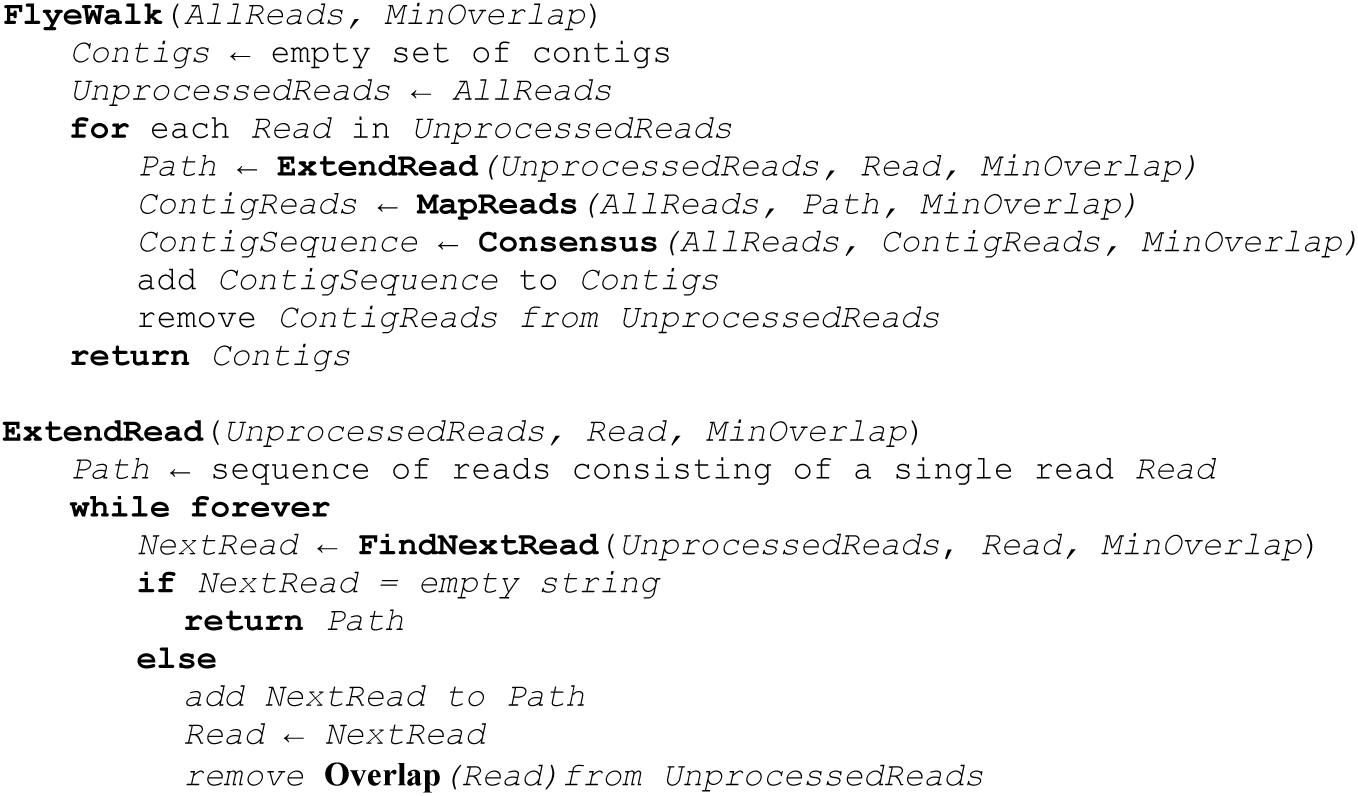
Pseudocode of the FlyeWalk algorithm. FlyeWalk iteratively extends each unprocessed read to form a path formed by the selected reads. Each such path contributes to a contig and **FlyeWalk** outputs the set of all contigs resulting from such path extensions. **ExtendRead** generates a random walk in the assembly graph, which starts at a given read and constructs a path (sequence of overlapping reads) that contributes to a constructed contig. It terminates when there are no unprocessed reads overlapping the current read by at least *MinOverlap* nucleotides. **FindNextRead** finds an unprocessed read that overlaps with the given read by at least *MinOverlap* nucleotides and returns an empty string if there are no such reads. **MapReads** finds all reads that map to a given path of reads (by at least *MinOverlap* nucleotides). **Consensus** constructs the consensus of all reads that contribute to a given contig. **Overlap** finds all reads that overlap a given read by at least *MinOverlap* nucleotides.

### Contracted assembly graph

Flye aligns all reads to the constructed assembly graph (Appendix “Aligning reads to assembly graph”) and classifies all edges in the assembly graph into unique and repeat edges (Appendix “Classifying edges of the assembly graph”). It further transforms the assembly graph into the *contracted assembly graph* by contracting all its repeat edges. Aligning a read to the assembly graph induces its alignment to the contracted assembly graph, and we focus on *spanning reads* that align to multiple edges in the contracted assembly graph. *Untangling* incident edges *e*=*(w,v)* and *f*=*(v,u)* in the contracted assembly graph amounts to substituting them by a single edge *(w,u).* Below we describe how Flye uses spanning reads to untangle the contracted assembly graph.

### Untangling the contracted assembly graph

A spanning read in the contracted assembly graph is classified as an *(e,f)-read* if it traverses two consecutive edges *e* and *f* in this graph. For each pair of incident edges *e* and *f* in the contracted assembly graph, we define *transition(e,f)* as the number of *(e,f)*-reads plus the number of *(f’,e’)-*reads, where *e’* and *f’* are *complementary* edges for *e* and *f*, i.e., edges representing a complementary strand.

Given a set of spanning reads in the contracted assembly graph, we construct a *transition graph* as follows. Each edge *e* in the contracted assembly graph corresponds to vertices *e*^*h*^ and *e*^*t*^ in the transition graph, representing the head and tail of *e*, respectively (a complementary edge for *e* correspond to the same vertices, but in the opposite order). Each (*e,f*)-spanning read defines an undirected edge between *e*^*h*^ and *f*^*t*^ in the transition graph with weight equal to *transition(e,f)*.

Note that the transition graph is bipartite for the simple case when the two subgraphs of the contracted assembly graphs corresponding to complementary strands do not share vertices. However, it is not necessarily bipartite in the case of genomes that contain long inverted repeats. Flye thus applies Edmonds’ algorithm (Edmonds, 1965) to find a maximum weight matching in the transition graph and uses this matching for untangling the contracted repeat graph. For each edge (*e*^*h*^,*f*^*t*^) in the constructed matching, Flye additionally checks the confidence of the transition between edges *e* and *f* (see Appendix “Additional details on untangling assembly graphs”) and untangles *e* and *f* for each edge (*e*^*h*^,*f*^*t*^) in the transition graph that passes this check. After iterative untangling of edges in the contracted assembly graph (and the corresponding iterative repeat resolution in the assembly graph), the assembly graph typically contains only long *unbridged* repeat edges that are not spanned by any reads.

**Figure 4.**
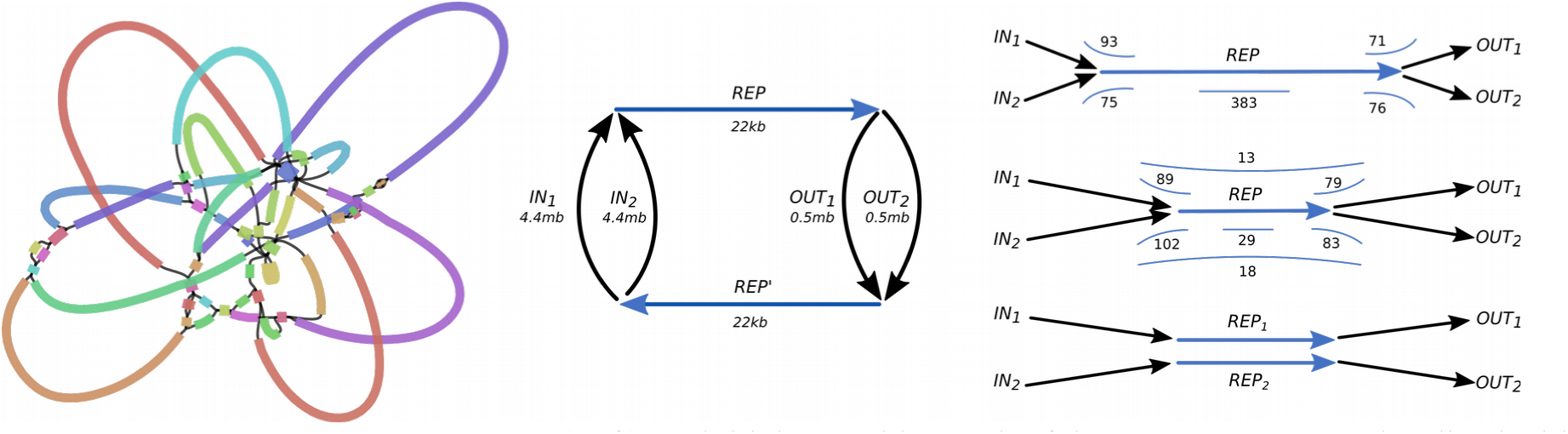
Resolving an unbridged repeat. (Left) An initial assembly graph of the EC9964 genome visualized with Bandage (Wick et al., 2015). (Middle) Assembly graph (after untangling repeats in the graph on the left using spanning reads) contains a single unbridged repeat *REP* (and its complement *REP’*) of length 22.0 kb. Two complementary strands are fused together in a single connected component. It is unclear whether the genome traverses the assembly graph as *IN_1_ REP OUT_1_ REP’* or as *IN_1_ REP OUT_2_ REP’.* (Right top) The edges ending in the initial node of the edge *REP* are denoted as *IN_1_* and *IN_2_*, and the edges ending in the terminal node of the edge *REP* are denoted *OUT_1_* and *OUT_2_*. 93, 71, 75, and 76 reads overlap with both *IN_1_* and *REP*, *IN_2_* and *REP*, *REP* and *OUT_1_* and *REP* and *OUT_2_*, respectively. The span of 383 reads falls entirely within the edge *REP*. (Right middle) After assigning 93 reads that overlap with both *IN1* and *REP* to the first repeat copy, and 71 reads that overlap with both *IN_2_* and *REP* to the second repeat copy, we “move forward” into the repeat and construct two differing consensus sequences for a prefix of *REP* of length 8.6 kb and divergence 9.8%. Similarly, we construct two consensus sequences for a 6.8 kb long suffix of *REP* with divergence 16.8% when we “move backward” into the repeat. The length of the repeat edge is reduced to 22.0 – 8.6 – 6.8 = 6.6 kb, resulting in the emergence of 13 + 18 = 31 spanning reads for this repeat, all of them supporting a *cis* transition (*IN_1_* with *OUT_1_* and *IN_2_* with *OUT_2_*). (Right bottom) Two resolved instances of the repeat with consensus sequences *REP_1_* and *REP*_2_ with divergence 6.9%.

### Resolving unbridged repeats

Resolving unbridged and nearly identical repeat instances is a difficult problem. SMS assemblers often fail to resolve unbridged repeats longer than 10 kb that are common even in bacterial genomes (Kamath et al., 2017). Although variations between different repeat copies allow for the possibility of resolving such repeats (at least in theory), no algorithm for resolving them using short reads has been successful so far. In the case of SMS reads, the problem is further complicated by high error rates that make it difficult to distinguish repeat copies with smaller divergence than 10%.

This challenge is somewhat related to the challenge of constructing phased diploid genome assemblies recently addressed by the developers of the Falcon and Canu assembler (Chin et al., 2016, Koren et al., 2017), but there are important differences specific to repeat resolution, notably, that repeats are flanked by unique edges. We leverage those differences in our approach, which we describe using the EC9964 dataset described in the Appendices “Resolving unbridged repeats” and “Information about the BACTERIA datasets”. For simplicity, we illustrate our algorithm using a 22.0 kb long unbridged repeat with two copies (referred to as R22) in the EC9964 genome. Flye and HINGE constructed topologically identical assembly graphs with this repeat unresolved (Figure 4, left), but Flye eventually resolved it using variations between its repeat copies.

Figure 4 shows this repeat with a corrupted consensus sequence *REP* in the subgraph of the assembly graph. Although it is impossible to resolve this repeat (i.e., to pair each entrance to this repeat with the corresponding exit) if its two copies are identical, it becomes possible if there are sufficient variations between these copies. The goal is to resolve the two distinct repeat copies and to transform the single sequence *REP* into two different repeat instances *REP_1_* and *REP_2_* as shown in Figure 4. Appendix “Resolving unbridged repeats” describes how Flye resolves unbridged repeats by (i) identifying the variations between repeat copies, (ii) matching each read with a specific repeat copy using these variations, and (iii) using these reads to derive a distinct consensus sequence for each repeat copy.

## RESULTS

### Datasets

We benchmarked various SMS assemblers using the following datasets:

- **BACTERIA dataset** contains 21 datasets from the National Collection of Type Cultures (NCTC), a project aimed at completing 3000 bacterial genomes using SMS technology (*http://www.sanger.ac.uk/resources/downloads/bacteria/nctc/*). We selected these 21 out of 997 datasets because they were studied in detail in Kamath et al. (2017) and used to benchmark various assemblers. Taking this into account, we only benchmarked Flye against HINGE on these datasets, since HINGE outperformed the other assemblers studied in Kamath et al. (2017).
- **YEAST dataset** contains Pacific Biosciences (PB) and Oxford Nanopore Technology (ONT) reads from *S. cerevisiae* S288c genome of length 12.1 Mb (16 chr.), used in the recent benchmarking of various SMS assemblers (Giordano et al., 2017). Similar to the original study, we used the full set of ONT reads (31x coverage) but down-sampled PB reads from the original 120x to 31x coverage to have their coverage and reads length distribution be similar to the ONT data.
- **WORM dataset** contains Pacific Biosciences reads (coverage 40X) from *C. elegans* genome of length 100 Mb (6 chr.) (*https://github.com/PacificBiosciences/DevNet/wiki/C.-elegans-data-set*).
- **HUMAN and HUMAN+ datasets.** The HUMAN dataset contain ONT reads (30x coverage) from the GM12878 human cell line (Jain et al., 2017). The HUMAN+ dataset combines the HUMAN-dataset with a dataset of ultra long ONT reads (5x coverage) generated using the recently developed ultra-long read protocol (Jain et al., 2017). The N50 is equal to 12.5 kb and 99.7 kb for the standard and the ultra-long reads, respectively.

### Benchmarking Flye, Canu, Falcon, HINGE, and Miniasm

We used QUAST version 5.0 (Mikheenko et al., 2018) to benchmark the latest versions of all assemblers (see Appendix “Additional information on benchmarking”). QUAST distinguishes between local and global misassemblies (such as inversions of large segments) using a parameter *minimum misassembly size.* However, using the default value for this parameter, tuned for short read assemblies, results in many local misassemblies with span below 6000 nucleotides reported by QUAST. These misassemblies often represent misalignments of long repetitive sequences such as the *Ty1* transposon in yeast. We thus decided to ignore local misassemblies by setting a large minimum misassembly size of 6,000 bp (the *minimum alignment identity* was set to a low 90%). Also, since Miniasm returns assemblies with a much larger number of mismatches and indels than other assemblers, it is not well suited for reference alignment-based quality evaluation, even with a low minimum alignment identity threshold. To make a fair comparison for Miniasm, we ran the ABruijn contig polishing module (Lin et al., 2016) on the Miniasm output to improve the accuracy of its contigs.

### Analyzing the BACTERIA dataset

Kamath et al., 2017 analyzed various NCTC datasets to demonstrate the advantages of the HINGE assembly graph approach. This analysis revealed that many NCTC datasets result in complex assembly graphs with long unbridged repeats that are nearly impossible to resolve. In contrast, various OLC assemblers attempting to resolve these repeats led to misassemblies.

We ignored small connected components in the assembly graphs (representing plasmids that do not share repeats with chromosomes) and classified assemblies of the BACTERIA datasets into three groups: (i) *complete* if the assembly graph consists of a single loop representing a circular chromosome, (ii) *semi-complete* if the assembly graph contains multiple edges but there exists a single solution of the *Chinese postman problem* (Edmonds and Johnson, 1973) in this graph, representing a complete circular chromosome, and (iii) *tangled* if the assembly graph is neither complete nor semi-complete. While HINGE does not distinguish between complete and semi-complete assemblies, we argue that ignoring this separation may lead to assembly errors. Indeed, a semi-complete assembly graph results in a unique assembly only in the case of uni-chromosomal genomes. In the case of multi-chromosomal genomes or in the case of plasmids that share repeats with a uni-chromosomal genome, there exist multiple possible assemblies from a semi-complete assembly graph. Since 10% of known bacterial species are multi-chromosomal (Jha et al., 2012) and since a large fraction of uni-chromosomal genomes have plasmids sharing repeats with the chromosome (Antipov et al., 2015), assuming that a semi-complete assembly graph results in a complete genome reconstruction is dangerous even in the case of bacterial genomes. Figure 5, left presents an example of a repeat that was resolved by HINGE based on the graph structure. Before resolving unbridged repeats, Flye assembled genomes from the BACTERIA dataset into four complete, one semi-complete, and 16 tangled assembly graphs. After resolving unbridged repeats, Flye assemblies resulted in six complete, three semi-complete, and 12 tangled assembly graphs. The number of edges in the tangled assembly graph constructed by Flye varied from 3 to 25.

**Figure 5.**
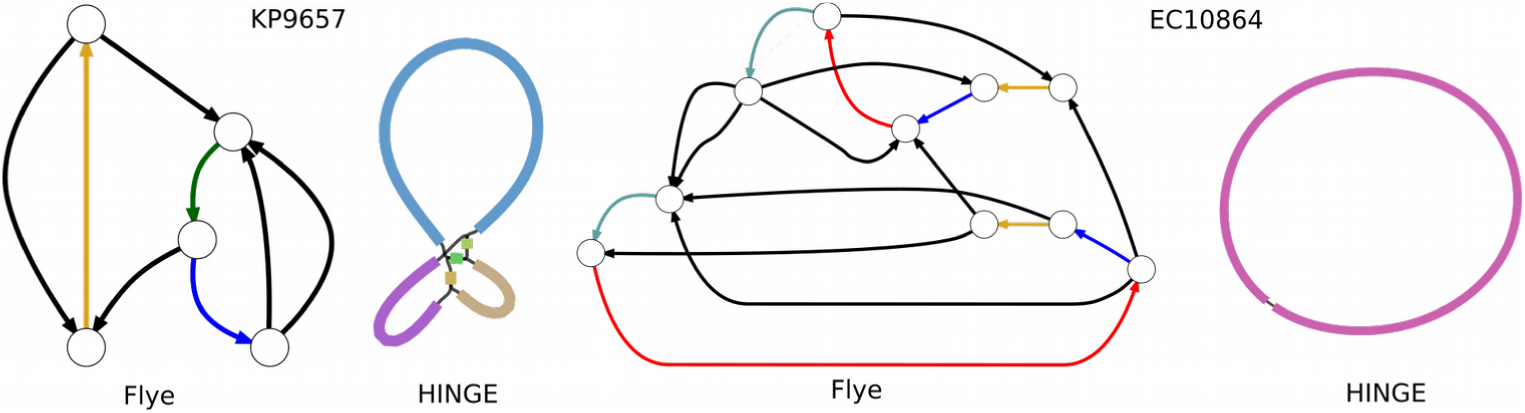
Differing assembly graphs constructed by Flye and HINGE. (Left) Flye and Hinge assembly graphs of the KP9657 dataset. There is a single unique edge entering into (and exiting) the unresolved “yellow” repeat and connecting it to the rest of the graph. Thus, this repeat can be resolved if one excludes a possibility that it is shared between a chromosome and a plasmid. In contrast to HINGE, Flye does not rule out this possibility and classifies the blue repeat as unresolved. (Right) Flye and Hinge assembly graphs of the EC10864 dataset show a mosaic repeat of multiplicity four formed by yellow, blue, red and green edges (the two copies of each edge represent complementary strands). HINGE reports a complete assembly into a single chromosome.

Flye disagrees with HINGE on only three BACTERIA datasets: (i) HINGE reports a complete assembly of KS11692, while Flye reports a tangled assembly graph that lacks circularization (the mean read length of the dataset is only 3928); (ii) HINGE reports a better assembly of the EC10864 dataset (Figure 5, right); (iii) Flye reports a better assembly of the EC9964 dataset after resolving an unbridged repeat (Figure 4). See Appendix “Information about the BACTERIA dataset” for more details.

### Analyzing the YEAST dataset

Table 1 illustrates that all tools but HINGE produced YEAST-PB assemblies with similar NGA50 ranging from 525 kb for Falcon to 567 kb for Canu (Flye fully assembled 11 out of 16 yeast chromosomes). Assembling this dataset with the original 120x coverage (rather than the downsampled 31x coverage) results in very similar assemblies, e.g., NGA50 increased from 561 kb to 581 kb for the Flye assembly. HINGE resulted in a lower NGA50 of 328 kb. Miniasm and Flye generated more accurate assemblies (10 misassemblies each) as compared to Canu, Falcon and HINGE (20, 26 and 25 misassemblies, respectively). Canu and Flye had the highest reference coverage (99.3% and 98.9%, respectively) and the most accurate contigs with the same average sequence identity 99.95%. Although Miniasm generated the least accurate contigs with only ≈90% identity with the reference, its contigs have become almost as accurate as the Flye contigs after the ABruijn-based polishing.

The YEAST-ONT assemblies show a similar trend, with Miniasm+ABruijn, Flye, and Canu producing the most accurate assemblies (8, 13 and 16 misassemblies, respectively). However, the Flye assembly was more contiguous (NGA50 = 769 kb versus 545 kb for Canu and 542 kb for Miniasm). Although most misassemblies represent small rearrangements, Miniasm and Falcon erroneously fused two chromosomes each in the YEAST-ONT dataset. For additional information, see Appendix “Assembly graph of the YEAST-ONT dataset”.

**Table 1.** Assembly statistics for the YEAST, WORM, and HUMAN/HUMAN+ datasets. Minimum misassembly size was set to 6000. Since Miniasm generates error-prone contigs, we polished them using ABruijn to provide a fair comparison.

| dataset | assembler | length (Mb) | #contigs | NG50 (kb) | reference coverage | reference % identity | misassemblies | NGA50 (kb) |
| --- | --- | --- | --- | --- | --- | --- | --- | --- |
| YEAST PB | Flye | 12.1 | 28 | **751** | **99.0%** | 99.95% | **10** | **561** |
|  | Canu | 12.2 | 43 | 588 | **99.3%** | 99.95% | 20 | **567** |
|  | Falcon | 11.9 | 42 | 562 | 96.7% | 99.81% | 26 | 525 |
|  | HINGE | 12.3 | 45 | 440 | 97.4% | 98.41% | 25 | 325 |
|  | Miniasm+ABrujn | 12.0 | 33 | 542 | 97.9% | 99.93% | **10** | 538 |
| YEAST ONT | Flye | 12.1 | 25 | **807** | **98.4%** | 99.04% | **13** | **773** |
|  | Canu | 12.0 | 29 | 774 | **98.6%** | 98.96% | 16 | 545 |
|  | Falcon | 11.9 | 41 | 716 | 97.4% | 98.81% | 18 | 637 |
|  | HINGE | 12.2 | 64 | 309 | 94.2% | 97.94% | 43 | 292 |
|  | Miniasm+ABrujn | 11.6 | 27 | 756 | 98.0% | 99.03% | **8** | 542 |
| WORM | Flye | 103 | 123 | 2,131 | **99.3%** | 99.93% | **62** | **1,682** |
|  | Canu | 108 | 262 | 2,264 | **99.2%** | 99.93% | 91 | 1,287 |
|  | Falcon | 101 | 106 | 2,291 | 98.7% | 99.78% | 89 | 1,286 |
|  | HINGE | 103 | 64 | **2,710** | 99.0% | 99.51% | 126 | 1,287 |
|  | Miniasm+ABrujn | 104 | 110 | 2,371 | 99.2% | 99.93% | 70 | **1,588** |
| HUMAN | Flye | 2,803 | 3,262 | 3,558 | **91.2%** | 98.50% | **587** | 3,194 |
|  | Canu | 2,646 | 2,883 | 3,103 | 88.5% | 97.20% | 748 | 2,244 |
|  | MaSuRCA | **2,877** | 8,773 | 4,670 | 90.5% | **99.79%** | 8,439 | **3,684** |
| HUMAN+ | Flye | 2,797 | 2,222 | 5,837 | **91.5%** | 98.50% | **939** | **4,474** |
|  | Canu | 2,759 | 2,327 | 5,824 | 89.0% | 97.19% | 1,288 | 3,778 |

### Analyzing the WORM dataset

Flye generated contigs with the lowest number of misassemblies (62), while other tools generated assemblies with the number of misassemblies varying from 70 for Miniasm+ABruijn to 126 for HINGE. Flye and Miniasm+ABruijn produced the largest NGA50 of 1,682 kb and 1,588 kb, respectively, while the other assemblies produced an NGA50 of 1,287 kb. Additionally, manual analysis of the reported misassemblies revealed the differences between the Flye assembly and the reference genome, which might be attributed to errors in the reference (see Appendix “Reconstruction of tandem repeats in the WORM dataset”). Miniasm+ABruijn, Flye, Canu, HINGE, and Falcon took 290, 480, 781, 803, and 945 minutes to assemble the WORM dataset (see Appendix “Additional details in benchmarking”).

### Analyzing the HUMAN and HUMAN+ datasets

We compared the Canu and Flye assemblies of the HUMAN and HUMAN+ datasets. We also downloaded a hybrid MaSuRCA assembly (Zimin et al., 2017) using the 30x ONT reads and the Illumina reads (available at the MaSuRCA web site).

For the HUMAN dataset, Flye and Canu generated assemblies with NGA50 equal to 3.2 Mb (587 assembly errors) and 2.2 Mb (748 assembly errors), respectively. Combining the HUMAN dataset of the ONT reads with short Illumina reads resulted in only a modest increase of NGA50 in the MaSuRCA hybrid assembly (3.6 Mb), while increasing the number of misassemblies by an order of magnitude (8,439 assembly errors). Manual inspection revealed that most errors in the MaSuRCA assembly were clustered within the difficult to assemble telomeric and centromeric chromosome regions. For the HUMAN+ dataset, Flye and Canu generated assemblies with NGA50 equal to 4.5 Mb (939 assembly errors) and 3.8 Mb (1,249 assembly errors), respectively. Based on a comparison of NGA50 statistics for Flye and MaSuRCA, it appears that when the coverage by long reads exceeds 30x, the contribution of short reads may be limited to reducing the error rate in the consensus sequences (for the ONT assemblies) rather than improving the quality of the assembly.

Flye, Canu, and MaSuRCA took 10,000, 40,000 and 50,000 CPU hours to generate assemblies of the HUMAN dataset (as reported by Canu, and MaSuRCA authors).

## DISCUSSION

In this manuscript we described the Flye algorithm for building the assembly graph of SMS reads and showed that repeat characterization is a useful complement for genome assembly algorithms. We showed that unbridged repeats that are not resolved with conventional read-spanning algorithms can be further untangled using the variations between repeat copies. Additionally, repeat graphs provide a useful framework for planning extra experiments for genome finishing.

We also compared Flye with popular long read assemblers. Flye and HINGE showed good agreement in the structure of the resulting assembly graphs on the BACTERIA datasets; however Flye significantly improved on HINGE on the more complex YEAST and WORM datasets. A possible explanation could be that Flye builds the assembly graph from accurate contigs while HINGE builds it from error-prone reads, thus limiting the graph resolution. Flye and Miniasm generated the most accurate and contiguous assemblies of the YEAST and WORM datasets compared to Canu, Falcon, and HINGE. Interestingly, both Miniasm and Flye (in contrast to Canu and Falcon) work with raw rather than error-corrected reads. In summary, Flye resulted in comparable or more accurate assemblies on all datasets. Specifically, in the case of WORM, HUMAN and HUMAN+ datasets, Flye generated significantly more contiguous and accurate assemblies than Canu.

## Acknowledgements

We are grateful to Bahar Beshaz and Sergey Nurk for their insightful comments.

## Availability

Flye is freely available at *http://github.com/fenderglass/Flye*. The supplementary files, including the assemblies generated by Flye are available at *https://doi.org/10.5281/zenodo.1143753*.

## APPENDICES

- Short read assemblers and the challenge of repeat resolution
- Consistent alignments
- Inconsistent alignments result in incorrect clustering-based repeat graph
- Flye constructs an accurate assembly graph from inaccurate FlyeWalk contigs
- Aligning reads to the assembly graph
- Classifying edges of the assembly graph
- Additional details on untangling assembly graphs
- Resolving unbridged repeats
- Defining the minimum divergence threshold
- Additional benchmarking information
- Information about the BACTERIA dataset
- Assembly graph of the YEAST-ONT dataset
- Reconstruction of tandem repeats in the WORM dataset

## Appendix: Short read assemblers and the challenge of repeat resolution

Although popular short read assemblers construct the assembly graph of *individual* reads (before resolving repeats using paired reads), they output a set of contigs (after the repeat resolution step) rather than an assembly graph constructed from the paired reads. For example, the SPAdes assembler (Bankevich et al., 2012) constructs the assembly graph of individual reads, uses it for repeat resolution, and outputs the resulting scaffolds (Prjibelsky et al., 2014). A better option would be to construct the assembly graph of these scaffolds (which are less tangled than the assembly graph of individual reads) and to apply the repeat resolution step again on this graph for a more accurate repeat resolution. Another advantage of this (less tangled) assembly graph lies in applications relating to hybrid assembly, e.g., co-assembly of short and long reads (Antipov et al., 2015, Wick et al., 2017). However, although some studies attempted to construct the assembly graph directly from paired reads (Vyahhi et al., 2012), the popular short read assemblers have failed to incorporate this approach into their pipelines so far.

## Appendix: Consistent alignments

Each multiple alignment of *m* sequences induces 
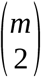
 pairwise alignments. A set of pairwise alignments (described by the repeat plot) is consistent if they can be combined into a single multiple alignment that induces each pairwise alignment in the set. However, the concept of multiple alignment is usually defined for the case of aligning multiple sequences rather than for aligning a sequence against itself. Below we describe the concept of a multiple self-alignment of a genome and define the notion of consistent pairwise self-alignments. This notion is important since A-Bruijn graphs result in a simple repeat graph in the case of consistent self-alignments but face various complications in the case of inconsistent self-alignments (see Pevzner et al., 2004 for examples of inconsistent alignments). The Flye algorithm bypasses the complications caused by inconsistent pairwise alignments enabling assemblies of long error-prone reads.

We define a multiple alignment of a single sequence against itself as simply a partition of its positions into non-overlapping subsets, with each subset corresponding to a column of the multiple alignment. For example, a multiple alignment of the sequence ACTGGCTGACT can be represented as a partition of 10 positions into six “painted” subsets: 
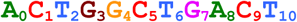
.
Figure A1 visualizes such a partitioning as a multiple alignment where each column represents positions from the same subset:

**Figure A1.**
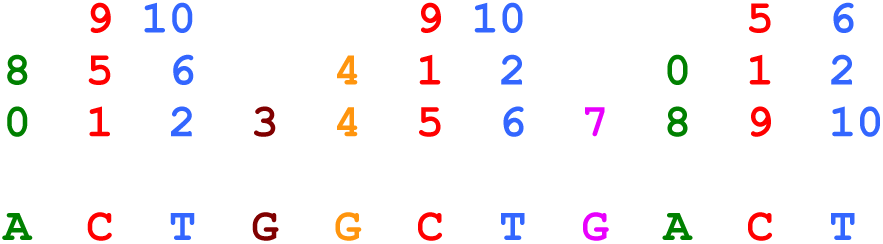
Multiple self-alignment defined by partitioning of the sequence 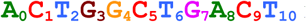 into six subsets. In difference from the traditional representation of a multiple alignment (where each entry represents a nucleotide in the multiple alignment matrix, the entries in the multiple self-alignment matrix represent positions in the sequence.

Every pair of numbers *i < j* in the same column of the multiple alignment defines a point (*i,j*) in the two dimensional plot. The collection of all such points defines the *dot plot* of the multiple alignment. We refer to a rectangle in the dot plot with lower left corner (*x,y*) and upper right corner (*x’,y’*) as (*x,y*, *x’,y’*). A local alignment between segments (*x,x’*) and (*y,y’*) of a genome defines a set of 2-dimensonal points in the rectangle (*x, y*, *x’, y’*) corresponding to matches in this alignment. A local alignment between segments (*x,x’*) and (*y,y’*) is *consistent* with the multiple alignment if the dot plot of the multiple alignment coincides with the dot-plot of the pairwise alignment within the rectangle (*x, y*, *x’, y’*). A set of local alignments is *consistent* if there exists a multiple alignment that is consistent with all local alignments in this set.

## Appendix: Inconsistent alignments result in incorrect clustering-based repeat graph

Figure A2 presents an example of an inconsistent alignment and illustrates that it results in a more complex repeat graph than the graph shown in Figure 2 in the main text. Although it may appear to be a minor annoyance in the case of the small example in Figure A2, inconsistent alignments may result in huge repeat graphs in the case of real genomes, making it difficult to analyze repeats in such cases. Also, although it may appear easy to add a missing diagonal (in the simple case shown in Figure A2) to make the alignment consistent, transforming inconsistent alignments into consistent ones is a challenging problem in the case of real genomes.

The punctilious repeat graph in Figure A2 has seven vertices (since there exist seven projections of alignment endpoints to the main diagonal), in contrast to the punctilious repeat graph in Figure 2 in the main text that has eight vertices. This deficiency of the punctilious repeat graph in Figure A2 motivates the algorithm for extending the set *Endpoints* as described in the main text. Note that the middle point of the long diagonal in Figure 2 represents an invalid point since only one of its projections belongs to the set of seven endpoint projections on the main diagonal. The algorithm described in the main text adds this projection to the set *Endpoints* and results in the same repeat graph as shown in Figure 2 in the main text.

**Figure A2.**
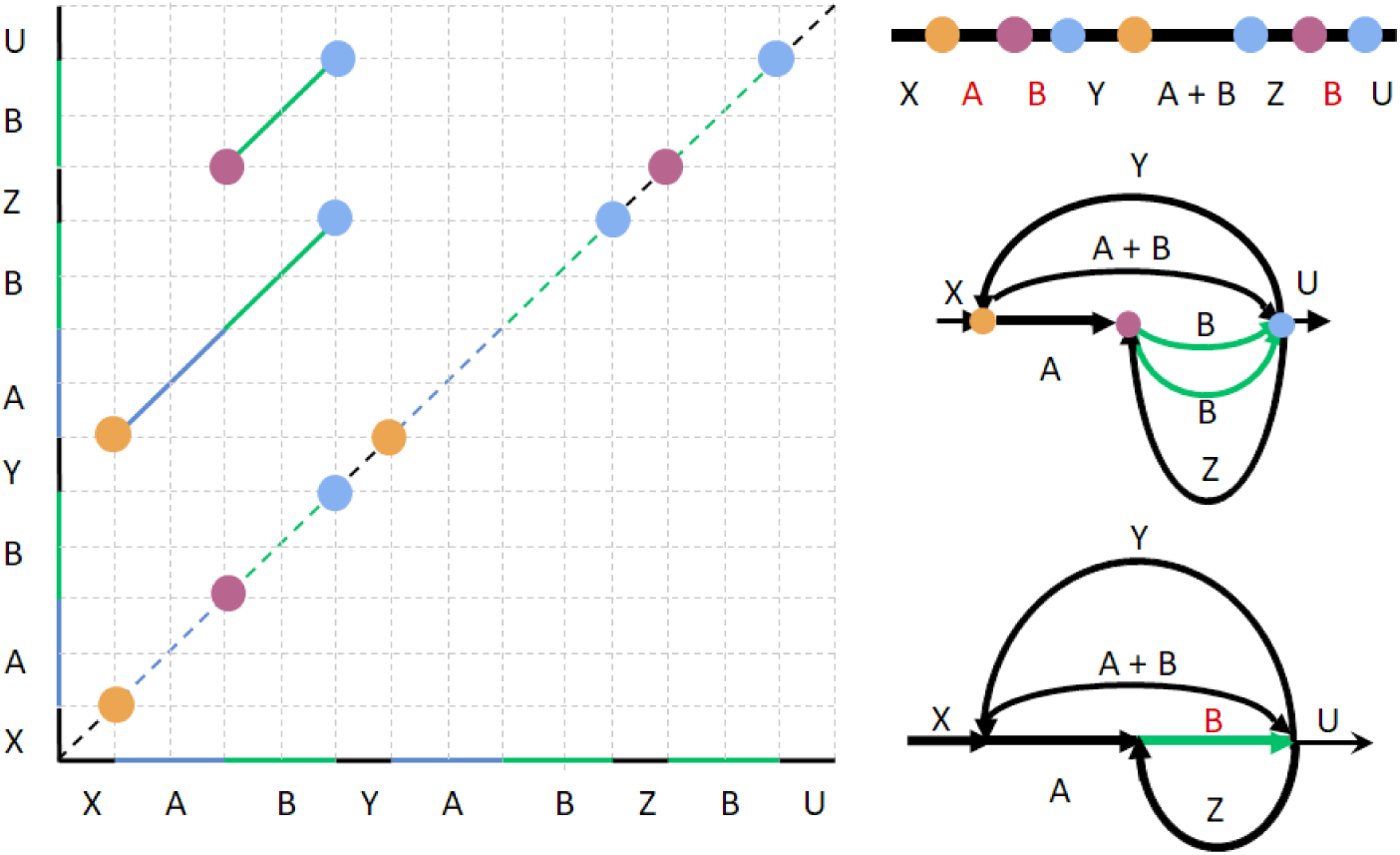
Inconsistent local alignments result in an “incorrect” repeat graph (as compared to the repeat graph shown in Figure 2 in the main text), thus necessitating an extension of the set of the alignment endpoints. (Left) Alignment paths for two local self-alignments within a genome XABYABZBU. Only two out of three pairwise alignments between instances of a mosaic repeat (AB, AB, and B) are shown since the third alignment did not pass the percent identity threshold, resulting in an inconsistent set of pairwise alignments. Alignment endpoints are clustered together if their projections on the main diagonal coincide or are close to each other (clusters of closely located endpoints for *d*=0 are painted with the same color). This clustering reveals three clusters with seven endpoints (Top right) Projections of the clustered endpoints on the main diagonal define seven vertices of the repeat graph. (Middle right). Gluing endpoints that belong to the same clusters. (Bottom right) Gluing parallel edges in the resulting graph (parallel edges are glued if there exists an alignment between their sequences), which results in the approximate repeat graph.

## Appendix: Flye constructs an accurate assembly graph from inaccurate FlyeWalk contigs

Consider the circular genomic tour in the assembly graph for the *E. coli* strain NCTC9964 formed by four edges IN1, REP, OUT1, REP’ and the corresponding complementary path formed by the complementary edges REP, OUT2, REP’, IN2 (Figure 4). Paths IN1, REP, OUT2, REP’ and IN2, REP, OUT1, REP’ form a set of incorrect contigs that however are assembled in the correct assembly graph by Flye. Below we explain why it holds for an arbitrary set of paths constructed by Flye from contigs constructed by `FlyeWalk`.

The construction of the repeat graph assumes that the genome *Genome* and the two-dimensional self-alignment matrix *Alignments* are given. Note that the alignment matrix defines the pairwise alignments (or lack of similarities) between any two substrings of the genome. Given a set of substrings from *Genome*, these alignments form *Alignments*-imposed pairwise alignments. A set of substrings of a genome is called a *covering set* if for every pair of consecutive positions in *Genome* there exists a substring containing these positions. In practice, we use a more stringent condition that requires that for each *m* consecutive positions in *Genome* (where *m* is a pre-defined threshold), there exists a substring (read) spanning all these positions.

A useful property of the repeat graph is that it can be constructed from alignments of substrings of *Genome* without knowledge of the entire sequence *Genome* and the entire matrix *Alignments*. As demonstrated in Pevzner et al., 2004, if reads form a covering set, then gluing of reads according to their *Alignments*-imposed similarities, produces the same repeat graph as gluing of the entire *Genome*. The order in which the reads are subjected to such gluing does not affect the resulting graph.

Let *Genome* be an (unknown) genomic sequence of an (unknown) length with an (unknown) alignment matrix *Alignments*. Let *s(1),…, s(t)* be a covering set of strings for *Genome*, and *A* be the collection of *t\**(*t* − 1)/2 sub-matrices of *Alignments* for every pair of substrings *s(i)* and *s(j)*. Every such 
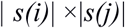
 sub-matrix is a snapshot of a “small” area of the matrix *Alignments*. The question is whether one can reconstruct the repeat graph from these snapshots rather than from the entire matrix *Alignments*.

We emphasize that coordinates of the strings *s(1),…, s(t)* and their ordering in the sequence *Genome* are unknown. However, Pevzner et al., 2004 demonstrated that the repeat graph of *Genome* coincides with the assembly graph of the sequence 
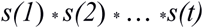
, the concatenate of all substrings (with delimiters) in any order. We use this result to demonstrate that the Flye assembly of inaccurate FlyeWalk contigs results in an accurate assembly graph.

Consider the set *Contigs* constructed by FlyeWalk and map all reads to all these contigs. Since FlyeWalk utilizes all reads, each read will be mapped to a single contig or to multiple contigs (for simplicity, we assume that chimeric reads have been removed). We now concatenate all reads starting from reads in the first contig, followed by reads in the second contig, etc., resulting in a sequence of reads:

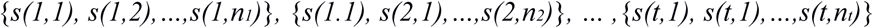

where *s(i,j)* stands for the *j*-th read in the *i-*th contig.

Since all reads are included in this concatenate, the repeat graph constructed from this concatenate coincides with the repeat graph of the genome (Pevzner et al., 2004). Note that gluing all reads from the first contig simply results in the consensus of this contig constructed by FlyeWalk. Iterating it for all contigs, we will get the following sequence of contigs as the result of these (but not all) gluing operations:

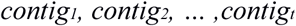

Therefore, the gluing of all contigs constructed by FlyeWalk (some of them may be misassembled) results in the same assembly graph as gluing all reads, and thus results in the repeat graph of the genome.

## Appendix: Aligning reads to the assembly graph

Flye aligns all reads to the constructed assembly graph using the concept of common *jump*-subpaths (Lin et al., 2016). First, each read is matched against the edges of the assembly graph. For each repeat edges in the assembly graph, we store all copies of the corresponding repeat (from the original contigs), rather than a single consensus of all sequences contributing to this repeat edge. We then match a read to *all* these copies and select the best alignment to improve the recruitment of reads to the edges of the assembly graph. If a read is mapped to multiple edges in the assembly graph, we find a maximum scoring path in the graph formed by such edges using dynamic programming.

## Appendix: Classifying edges of the assembly graph

After constructing the assembly graph, Flye aligns all reads to this graph and forms a *read-path* for each read. Given alignments of all reads against the assembly graph, Flye computes the mean depth of coverage *cov* across the entire assembly graph and classifies an edge as *low-coverage* (if its coverage is below 2\**cov*) and *high-coverage* otherwise. While most low coverage edges are unique (traversed only once in the genomic tour), some of them are repetitive since the coverage varies along the genome.

To accurately classify unique and repetitive edges in the assembly graph, Flye complements coverage-based classification using information about the read-paths. An edge *e’* in the assembly graph is a *successor* of an edge *e* if it follows *e* in one of the read-paths. A low-coverage edge is classified as *unique* if it has a single successor. All other edges (i.e., high-coverage edges or low-coverage edges with multiple successors) are classified as repetitive. To avoid classifying chimeric connection in the assembly graph as successor edges and to minimize the influence of misaligned reads, Flye imposes an additional restriction on the edge to classify it as a successor: a fraction of the reads supporting each successor (among all reads contributing to the successor of a given edge) should exceed a threshold.

## Appendix: Additional details on untangling assembly graphs

After reconstructing the maximum weight matching, which defines the set of edges in the transition graph, Flye additionally checks each of the inferred edges as follows. For each edge *(u, v)* from the matching, we compute the total weight *s* of all edges in the transition graph adjacent to *u* or *v*. If *transition(u, v)* < *s* / 2, the edge is called *weak* and is consequently ignored. Weak edges typically occur because long repeats may be spanned by a few reads in an ambiguous way.

Flye iteratively untangle edges and finds maximum weight matchings until no extra repeats can be resolved. Note that some edges that were initially classified as repetitive may become unique during the next iteration of the algorithm (for example, if they were a part of a bigger mosaic repeat that was partially resolved).

## Appendix: Resolving unbridged repeats

Flye takes advantage of the small variations between different repeat copies to resolve unbridged repeats. The basic idea is to (i) identify the variations between repeat copies, (ii) match each read with a specific repeat copy using these variations, and (iii) use these reads to derive a distinct consensus sequence for each repeat copy. The success of this approach is contingent upon the presence of a sufficient number of variations between the different repeat copies. Therefore, the first step is to estimate the number and positions of variations between the repeat copies and to calculate the *divergence* of the various repeat copies from reads alone.

**Revealing variable positions within repeats.** To reveal the variable positions within a repeat, we map all reads to the consensus sequence of the repeat (constructed by the ABruijn polishing procedure) and generate a multiple sequence alignment of all reads that are contained within or overlap with the repeat. Afterwards, we determine the most frequent symbol in each column of the alignment, count the occurrences of every other symbol (A, C, G, T, or “-”) in this column, and determine the substitution, deletion, and insertion rate for each symbol in each column as described in Lin et al., 2016. If the substitution, deletion, or insertion rate for a symbol is higher than a predetermined threshold, the position is called a *tentative divergent position*. The *repeat divergence* is then estimated by dividing the total number of tentative divergent positions by the length of the repeat.

Since substitutions and indels frequently occur in non-divergent positions of SMS reads, it is important to choose thresholds that are higher than this background distribution of errors, but low enough to capture the truly divergent positions. To select thresholds, we analyzed the R22 repeat in the assembly graph of the EC9964 dataset (Figure 4). Since this repeat has many variations between two repeat copies (1515 (6.9%) true divergent positions, (943 substitutions (4.3%), 346 deletions (1.6%), and 226 insertions (1.0%)), it was manually resolved with high confidence. With the accurate sequences of two repeat copies identified, we are able to distinguish the background distribution of mutation rates imposed by errors in non-variable positions from the mutation rates at variable positions to identify appropriate thresholds.

We mapped reads from EC9964 to the consensus of the R22 repeat and calculated the substitution, deletion, and insertion rates for the second most frequent symbol at each column of the alignment. Figure A3 illustrates that variable positions feature higher substitution, deletion, and insertion rates than non-variable positions within a repeat. We thus identify tentative divergent positions based solely on mutation rates by selecting a mutation rate threshold that provides a good separation between the two distributions (0.1, 0.2, and 0.3 for substitutions, deletions, and insertions, respectively). This results in the identification of 924 out of 943 substitutions, 270 out of 346 deletions and 54 out of 226 insertions for the R22 repeat. At the same time, we misclassified 81 identical positions as divergent (61 substitutions, 5 deletions, and 15 insertions), resulting in a false positive rate of 0.4%. In all, we identified 1329 tentative divergent positions, which leads to a divergence estimate of 6.0%, a slight underestimation of the true divergence rate of 6.9%.

**Figure A3.**
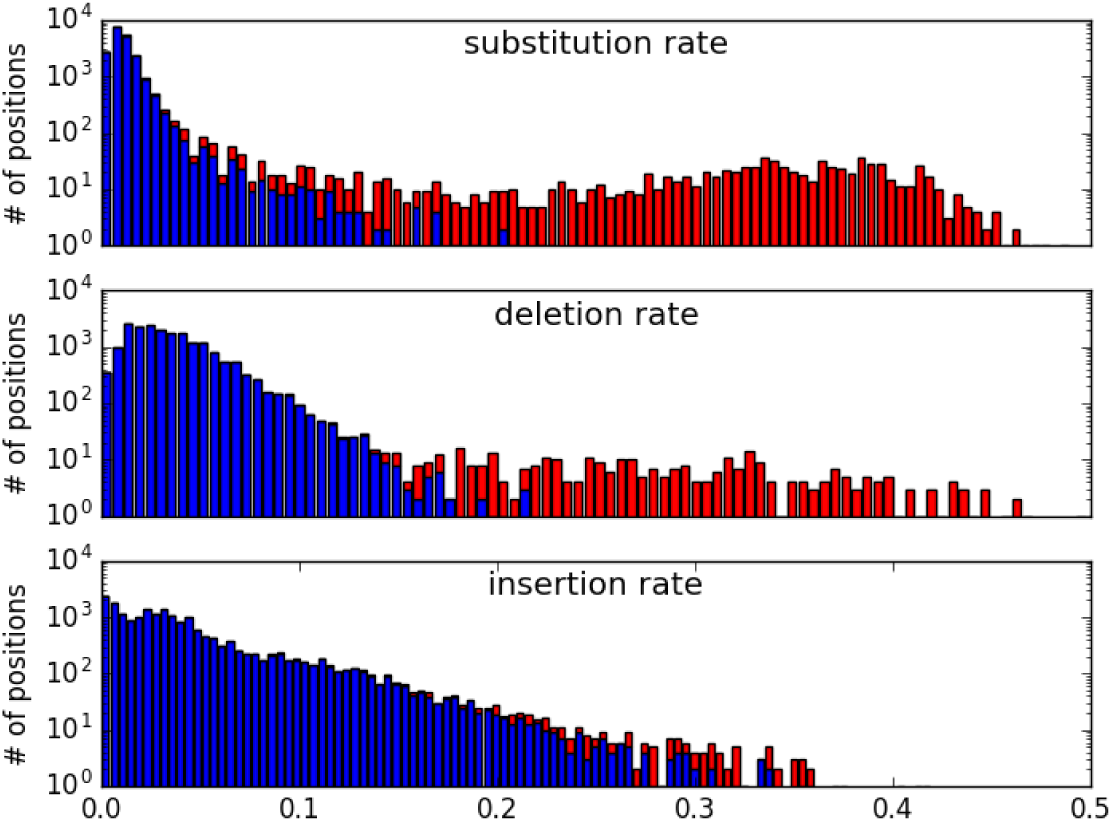
Separating divergent and identical positions within repeats using mutation rates computed for R22 repeat in the EC9964 dataset. Substitution (top), deletion (middle), and insertion (bottom) rates for the second most frequent symbol at each position in the multiple alignment of reads. Mutation rates for identical (divergent) positions are shown in blue (red). The number of positions with a given mutation rate (*y*-axis) is shown in a logarithmic scale. The cutoffs 0.1, 0.2, and 0.3 result in a good separation of identical and divergent positions for substitutions, deletions, and insertions, respectively.

**Resolvable and unresolvable repeats.** Based on the identified tentative divergent positions, we classify a repeat as resolvable based on the following two criteria:

- *The divergence rate exceeds a minimum divergence threshold.* Based on simulated data, we set up a minimum 0.1% divergence threshold, i.e., at least one divergent position per each 1000 bp on average (see Appendix “Defining the minimum divergence threshold”). When the divergence rate falls below 0.1%, there is often a shortage of reads covering multiple divergent positions, which is necessary for successful repeat resolution.
- *The maximal distance between consecutive divergent positions does not exceed the maximum distance threshold.* E.g., if two consecutive divergent positions are 15 kb apart but the maximal read length is 10 kb, there will be no reads spanning these positions that can be used for repeat resolution. Moreover, it turns out that the maximal read length is too conservative of a threshold, since consecutive divergent positions may be less than that length but the repeat is still unresolvable (Figure A4). On the other hand, we found that selecting the average read length as the threshold is too lenient. Based on our analysis of the BACTERIA dataset, we set the default threshold for the maximal distance between consecutive divergent positions as twice the average read length, which varies from 12 kb to 20 kb in the BACTERIA datasets.

If either of the above criteria does not hold, the repeat is classified as unresolvable. Note that some repeats initially classified as resolvable may be later classified as *unresolved* if the algorithm described below fails to resolve them.

**Figure A4.**
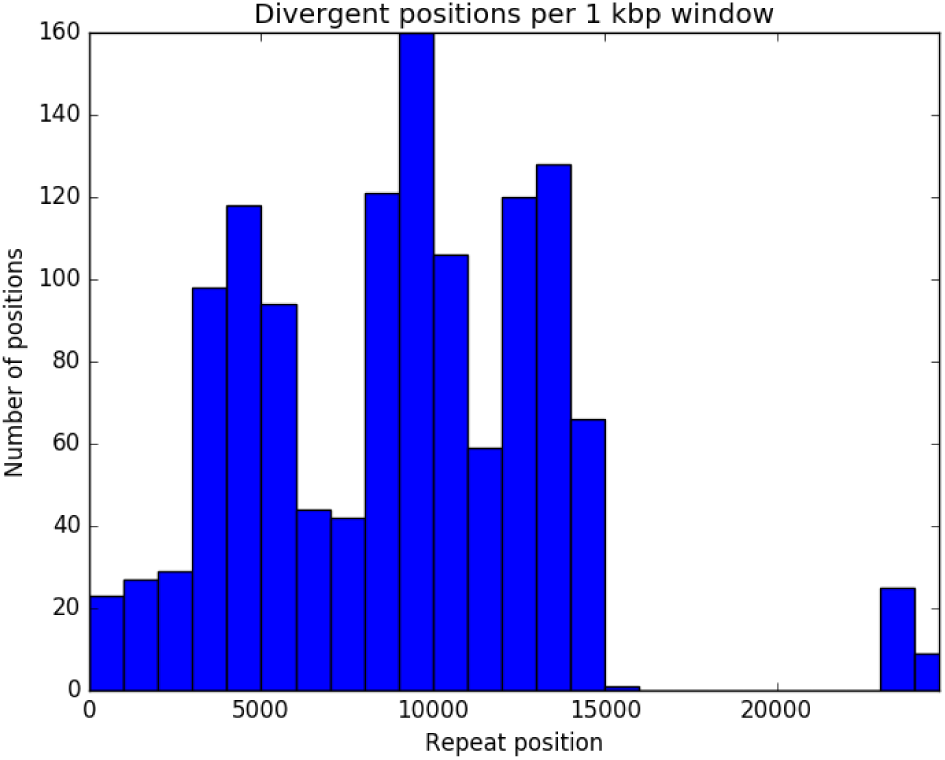
Histogram of divergent positions for a 24 kb long repeat from the EC9007 dataset with. a large 8 kb gap between two consecutive divergent positions located at 15002 and 23150 nucleotides from the start of the repeat. The repeat remained unresolved since there is only one read spanning this gap, which does not reveal a confident pairing of the incoming and outgoing edges for this repeat.

**Repeat Resolution Algorithm.** The idea of the algorithm is to utilize the tentative divergent positions to assign each read to a specific repeat copy and then use the assigned reads to derive a distinct consensus sequence for each repeat copy. For example, the 93 reads that span edges *IN*_1_ and *REP* (Figure 4 in the main text) can be assigned to one repeat copy and the 71 reads that span edges *IN*_2_ and *REP* can be assigned to another repeat copy. However, it is unclear how to assign other reads overlapping with the edge *REP* to a specific repeat copy.

In each iteration of the algorithm, reads are assigned to a specific repeat copy and then all the reads assigned to each repeat copy are used to construct a consensus sequence for that copy. Thus, as the algorithm proceeds, more and more reads are assigned to specific repeat copies and the consensus sequence for each repeat copy grows longer and longer. The algorithm terminates when no new reads can be assigned to read copies and the consensus sequences stop growing in length. There are two goals: to obtain distinct consensus sequences for each repeat copy and to determine the correct pairings of incoming and outgoing edges for each repeat copy.

Initially, the algorithm assigns all reads spanning edges *IN*_1_ and *REP* (*IN*_2_ and *REP*) to the first (second) repeat copy and computes the consensus of each repeat copy using the recruited reads. Since the recruited reads do not span the entire edge *REP*, we only construct two consensus sequences corresponding to prefixes of *REP*, where there is substantial read coverage by the recruited reads (at least 10X for each repeat copy to ensure that consensus sequences are sufficiently accurate). Both consensus sequences are truncated to the length of the shortest consensus sequence to prevent bias in the read recruitment process in future iterations. In the case of the EC9964 dataset, we constructed two consensus sequences corresponding to 8.6 kb long prefixes of *REP* with divergence 9.8% (Figure 4). As a result, we now have two consensus sequences for the entire edge *REP* that differ in some of the first 8.6 kb but coincide in the remaining part.

The two constructed consensus sequences serve as two templates for mapping reads to specific repeat copies in successive iterations. In this way, we can be conservative in gradually constructing the consensus sequences from only reads that have been assigned to a specific repeat copy with high confidence. In each successive iteration, the algorithm follows a series of three steps:

- *Evaluating the tentative divergent positions.* We map all classified reads again, this time to two consensus copies of the repeat (rather than a single consensus copy as in the initial iteration) to construct a more accurate alignment. We further consider the consensus sequence of each repeat copy and compare the most frequent symbols (A, C, G, T, “-”) occurring in the set of already classified reads for each repeat copy at each tentative divergent position. If the most frequent symbol at a position differs for two repeat copies, then that position is called a *confirmed divergent position*, otherwise it is called a *rejected divergent position*. The most frequent symbols of all the confirmed divergent positions for a certain repeat copy represent a “signature” of this copy. Since some positions within a repeat may not have been reached by the two consensus sequences yet, they remain classified as tentative divergent positions.
- *Assigning reads to various repeat copies.* We now map all unclassified reads to two consensus copies of the repeat and utilize the confirmed divergent positions to assign some unclassified reads to a specific repeat copy. For each read, we compute its *vote* for each repeat copy as the number of confirmed divergent positions at which the symbol of the read agrees with the consensus of this repeat copy. The read is assigned to a specific repeat copy if its vote for this copy is larger than the vote for another copy (the read remains unassigned in the case of ties).
- *Constructing new consensus sequences for each repeat copy.* We use all reads that have been assigned to a specific repeat copy to construct a new consensus sequence of this copy. The consensus is only called up to where the coverage of the reads is at least 10X in both repeat copies to ensure that consensus sequences are accurate, and then both consensus sequences are truncated to the length of the shortest consensus sequence. The algorithm then proceeds to the next iteration unless no new reads mapping to the original repeat consensus were classified or all of the consensus sequences are identical to those in the previous iteration, for which cases it terminates.

At the conclusion of the algorithm, a consensus sequence has been constructed for each repeat copy and a set of confirmed divergent positions for each repeat copy has been obtained.

Although we discussed the algorithm as “moving forward” into the repeat” (e.g., moving ahead from edges *IN*_1_ and *IN*_2_ in Figure 4 in the main text), the same procedure is performed by “moving backward” in the opposite direction (e.g., moving backwards from edges *OUT*_1_ and *OUT*_2_ in Figure 4 in the main text), or equivalently, moving forward along the reverse complement of the repeat. Eventually, as the repeat consensus sequences extend forward and backward (Figure 4 in the main text), this procedure may result in the emergence of *spanning reads,* i.e., reads that are assigned to both a repeat copy originating from one of the incoming edges (*IN*_1_ or *IN*_2_) and a repeat copy originating from one of the outgoing edges (*OUT*_1_ or *OUT*_2_). Spanning reads are grouped depending on which incoming/outgoing edges they are assigned to: *IN*_1_ and *OUT*_1_, *IN*_2_ and *OUT*_2_, *IN*_1_ and *OUT*_2_, and *IN*_2_ and *OUT*_1_. We further classify all spanning reads into one of two categories called *cis* (IN_1_/OUT_1_ and IN_2_/OUT_2_) and *trans* (IN_1_/OUT_2_ and IN_2_/OUT_1_) since there are only two ways to resolve the repeat: pairing IN_1_/OUT_1_ with IN_2_/OUT_2_, or pairing IN_1_/OUT_2_ with IN_2_/OUT_1_.

When the number of spanning reads in one of the categories exceeds a threshold (the default value is five) and exceeds the number of spanning reads in another category by at least a factor of two, all reads in the “winning” category are assigned to the corresponding repeat copies and the consensus of each repeat copy is re-computed. Afterwards, Flye continues moving forward and backward into the repeat recruiting more and more spanning reads until all remaining reads are classified as spanning reads.

If our attempts to resolve the repeat did not result in the emergence of spanning reads or if the conditions above on the number of spanning reads do not hold, the repeat is classified as unresolved (note that some resolvable repeats may be classified as unresolved). Note that even in the case of unresolved repeats, our algorithm still generates more accurate consensus sequences for the prefixes and suffixes of the repeat. Table A1 summarizes the results of the algorithm on 11 datasets from BACTERIA containing repeats of multiplicity two (10 datasets from BACTERIA do not contain such repeats).

**Table A1.** Analysis of unbridged repeats of multiplicity 2 in genomes from the BACTERIA dataset. The results of repeat resolution after running Flye for the 11 out of 21 BACTERIA datasets that contain repeats of multiplicity 2. The label of each dataset denotes the bacterial species, its strain, and a letter referring to a specific repeat of multiplicity 2 found in the assembly graph (e.g. EC5052A, EC5052B, and EC5052C refer to 3 repeats A, B, and C present in the assembly graph for the *E. coli* NCTC5052 dataset). Bolded rows refer to repeats that were successfully resolved using Flye. The divergence is computed by dividing the number of tentative divergent positions (DP) by the repeat length. Max. DP distance refers to the maximal distance between tentative divergent positions in a repeat. Remaining gap refers to the length of the repeat remaining without separate consensus sequences for each copy after we have “moved into the repeat” from both the forward and reverse directions. Spanning reads refer to reads that are assigned to both a repeat copy originating from an incoming edge and a repeat copy originating from an outgoing edge (as defined in the text). *cis* spanning reads are assigned to either *IN_1_* and *OUT_1_* or *IN_2_* and *OUT_2_*, and *trans* spanning reads are assigned to either *IN_1_* and *OUT_2_* or *IN_2_* and *OUT_1_*.

| Dataset | Repeat length (kb) | Coverage | Divergence | #Tentative DP | #Confirmed DP | Max. DP distance | Remaining gap (kb) | #cis spanning reads | #trans spanning reads |
| --- | --- | --- | --- | --- | --- | --- | --- | --- | --- |
| KN5052A | 32 | 131X | 4.8% | 1540 | 623 | 6952 | 1k | 0 | 1 |
| KN5052B | 39 | 231X | 0.3% | 150 | 0 | 7122 | 2k | 0 | 0 |
| **KN5052C** | **32** | **174X** | **1.3%** | **410** | **246** | **4487** | **0** | **31** | **40** |
| **EC7921A** | **12** | **177X** | **17%** | **2085** | **1213** | **367** | **0** | **26** | **335** |
| EC9006A | 22 | 140X | 0.5% | 138 | 87 | 4395 | 4k | 0 | 1 |
| EC9007A | 24 | 329X | 5.4% | 1449 | 70 | 8148 | 6k | 1 | 0 |
| **EC9012A** | **30** | **106X** | **4.9%** | **1482** | **1184** | **1658** | **0** | **636** | **21** |
| EC9016A | 16 | 95X | 8.2% | 163 | 88 | 2768 | 1k | 0 | 0 |
| KN9657A | 36 | 85X | 0.1% | 48 | 0 | 6006 | 26k | 0 | 0 |
| **EC9964A** | **22** | **138X** | **6.0%** | **1329** | **767** | **580** | **0** | **666** | **0** |
| **EC11022A** | **30** | **83X** | **5.9%** | **1784** | **988** | **2236** | **0** | **25** | **73** |
| EC11022B | 24 | 82X | 0.2% | 61 | 13 | 2754 | 12k | 0 | 0 |
| **SA11962A** | **8** | **292X** | **9.4%** | **779** | **607** | **291** | **0** | **691** | **0** |
| KL12158B | 12 | 88X | 0.1% | 14 | 0 | 4795 | 5k | 0 | 0 |

## Appendix: Defining the minimum divergence threshold

To determine the minimum divergence threshold for which the repeat resolution algorithm could be applied successfully, we simulated several repeats of multiplicity two of length 10 kb, 20 kb, and 40 kb, with divergence rates ranging from 0.01% to 0.45%. Variations between the different copies of these repeats were introduced by adding substitutions and indels randomly to both copies until the desired divergence rate was reached. Next, we simulated Pacific Biosciences reads from these repeats with coverage 100X, mean error rates of 15% and read lengths between 5 kb and 15 kb. When the repeat resolution algorithm was applied to these datasets, we found that all simulated repeats with divergence rate greater than 0.1% were successfully resolved. We thus chose 0.1% to be the minimum divergence threshold.

## Appendix: Additional details on benchmarking

**Software versions.** We used the latest versions of all assemblers that were available in October 2017. However, we ran Canu 1.3 instead of the latest Canu 1.6 version on the YEAST-ONT dataset since version 1.6 failed to produce output within 24 hours. All assemblers were benchmarked with their default/suggested parameters:

- Flye - 2.3 (commit e73884c)
- Canu - 1.6 (commit 2d3f55a)
- Falcon - 0.3.0 (FALCON-Integrate commit 7498ef9)
- HINGE - 0.5.0 (commit 79fdf66)
- Miniasm - 0.2-r168-dirty (commit 40ec280) / Minimap 0.2-r124-dirty (commit fdf32ad)
- QUAST 5.0-dev (commit 974846a)

**Table A2.** Running time and memory usage of various SMS assemblers. We used a desktop machine with Intel(R) Core(TM) i7-4790 CPU @ 3.60GHz (up to 8 threads available) for the YEAST dataset assemblies; a single computational node with Intel(R) Xeon(R) CPU X5680 @ 3.33GHz for the WORM dataset assemblies (up to 24 threads available). Since we performed an additional polishing step on the Miniasm output, the running time for Flye and Miniasm are given for runs with and without contig polishing.

| Dataset | Assembler | Wall clock time | Peak memory usage |
| --- | --- | --- | --- |
| YEAST PB<br>12 Mb genome, 31x<br>8 threads max | Flye (w/o polishing) | 25m (9m) | 7G (7G) |
|  | Canu | 93m | 5G |
|  | Falcon | 62m | 10G |
|  | HINGE | 9m | 5G |
|  | Miniasm+ABruijn (Miniasm) | 16m (1m) | 5G (5G) |
| YEAST ONT<br>12 Mb genome, 31x<br>8 threads max | Flye (w/o polishing) | 28m (18m) | 7G (7G) |
|  | Canu | 283m | 8G |
|  | Falcon | 103m | 11G |
|  | HINGE | 11m | 8G |
|  | Miniasm+ABruijn (Miniasm) | 31m (3m) | 5G (5G) |
| WORM<br>100 Mb genome, 40x<br>24 threads max | Flye (w/o polishing) | 480m (348m) | 27G (27G) |
|  | Canu | 781m | 20G |
|  | Falcon | 945m | 18G |
|  | HINGE | 803m | 52G |
|  | Miniasm+ABruijn (Miniasm) | 290m (10m) | 23G (23G) |

## Appendix: Information about the BACTERIA dataset

**Table A3.** Comparison of Flye and HINGE on bacterial genomes from the BACTERIA dataset. HINGE results were taken from (Kamath et al., 2017). Since most sub-datasets in the BACTERIA dataset were generated with an older P5-C3 chemistry, the *MinOverlap* parameter of Flye was set to 3000 by default to account for shorter mean read lengths. Additionally, for the EC4450, EC8333 and KC12993 datasets, we used *MinOverlap* = 2000 to account for even shorter mean read lengths. EC9016 dataset was run using *MinOverlap* = 5000, as the lower parameter values produced fragmented assemblies. “Tangled*” stands for “Tangled/Lack Circularization”. “n/a” indicates that the assembly graph has no unbridged repeats of multiplicity two.

| Dataset | Bacterial species | Flye | Flye + unbridged repeat resolution | HINGE | Agreement |
| --- | --- | --- | --- | --- | --- |
| EC4450 | *Escherichia coli* | Tangled * | n/a | Tangled | Yes |
| KP5052 | *Klebsiella pneumoniae* | Tangled | Tangled | Tangled | Yes |
| SA6134 | *Staphylococcus aureus* | Complete | n/a | Complete | Yes |
| EC7921 | *Escherichia coli* | Tangled | Complete | Complete | Yes |
| EC8333 | *Escherichia coli* | Tangled * | n/a | Tangled | Yes |
| EC8781 | *Escherichia coli* | Tangled | n/a | Tangled | Yes |
| EC9002 | *Escherichia coli* | Complete | n/a | Complete | Yes |
| EC9006 | *Escherichia coli* | Tangled | Tangled | Tangled | Yes |
| EC9007 | *Escherichia coli* | Tangled | Tangled | Tangled | Yes |
| EC9012 | *Escherichia coli* | Tangled | Semi-complete | Complete | Yes |
| EC9016 | *Escherichia coli* | Tangled | Tangled | Tangled | Yes |
| EC9024 | *Escherichia coli* | Tangled | n/a | Tangled | Yes |
| EC9103 | *Escherichia coli* | Complete | n/a | Complete | Yes |
| KP9657 | *Klebsiella pneumoniae* | Tangled | Tangled | Tangled | Yes |
| EC9664 | *Escherichia coli* | Tangled | Complete | Tangled | No |
| EC10864 | *Escherichia coli* | Tangled | n/a | Complete | No |
| EC11022 | *Escherichia coli* | Tangled | Semi-complete | Complete | Yes |
| KS11692 | *Klebsiella sp* | Tangled* | n/a | Complete | No |
| KP12158 | *Klebsiella planticola* | Semi-complete | Semi-complete | Complete | Yes |
| KC12993 | *Kluyvera cryocrescens* | Complete | n/a | Complete | Yes |

## Appendix: Assembly graph of the YEAST-ONT dataset

**Figure A5.**
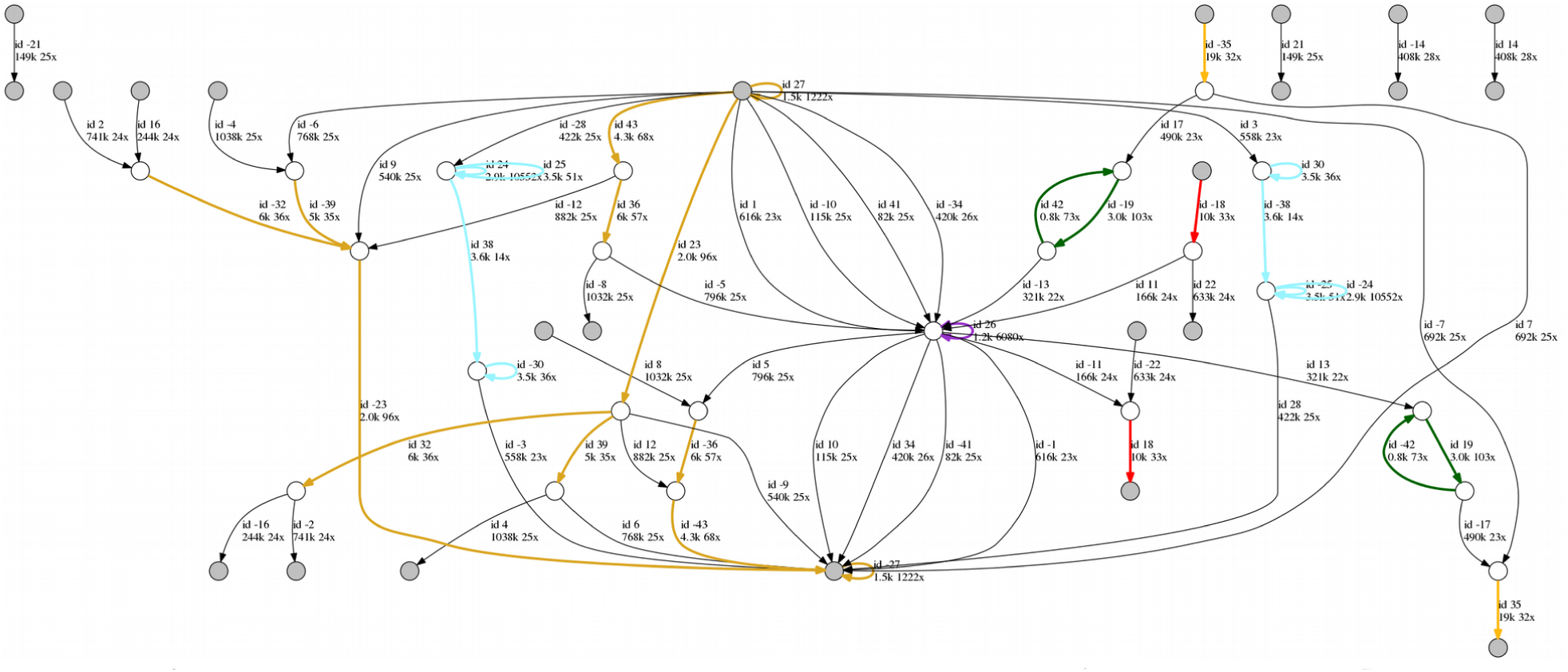
The Flye assembly graph of the YEAST-ONT dataset. Flye assembled the YEAST-ONT dataset into a graph with 22 unique and 14 repeat edges and generated 23 contigs as unambiguous paths in the assembly graph. Each unique contig is formed by a single unique edge and possibly multiple repeat edges, while repetitive contigs consist of the repetitive edges which were not covered by the unique contigs. The assembly graph was generated using the graphviz tool.

## Appendix: Reconstruction of tandem repeats in the WORM dataset

**Figure A6.**
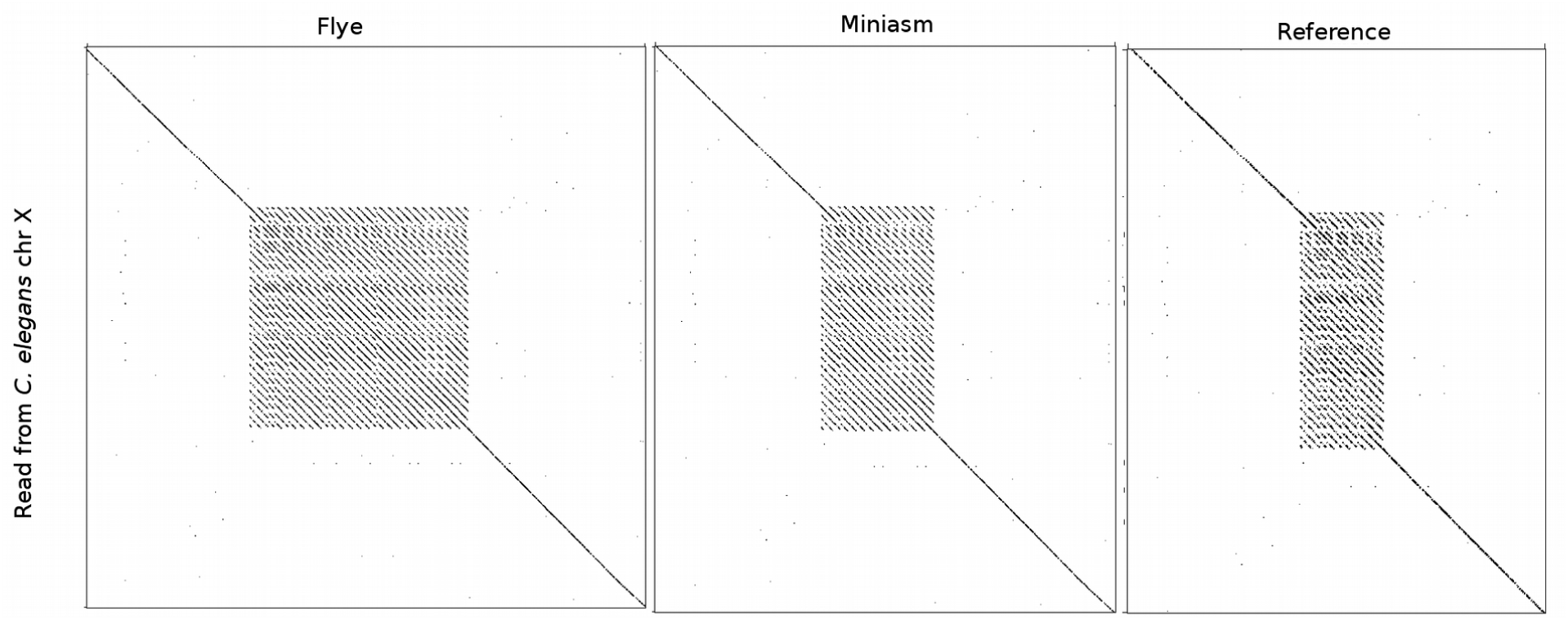
(Case A) Dot-plots showing the alignment of a read from chromosome X against the Flye assembly (left), the Miniasm assembly (middle) and the reference genome of *C. elegans* (right). The reference genome contains a tandem repeat of length 1.9 kb (10 copies) on chromosome X. In contrast, the Flye and Miniasm assemblies of this region suggest a tandem repeat of length 5.5 kb (27 copies) and 2.8 kb (13 copies), respectively. 15 reads that spans over the tandem repeat supports the Flye assembly (the mean length between the flanking unique sequence matches the cluster length reconstructed by Flye).

**Figure A7.**
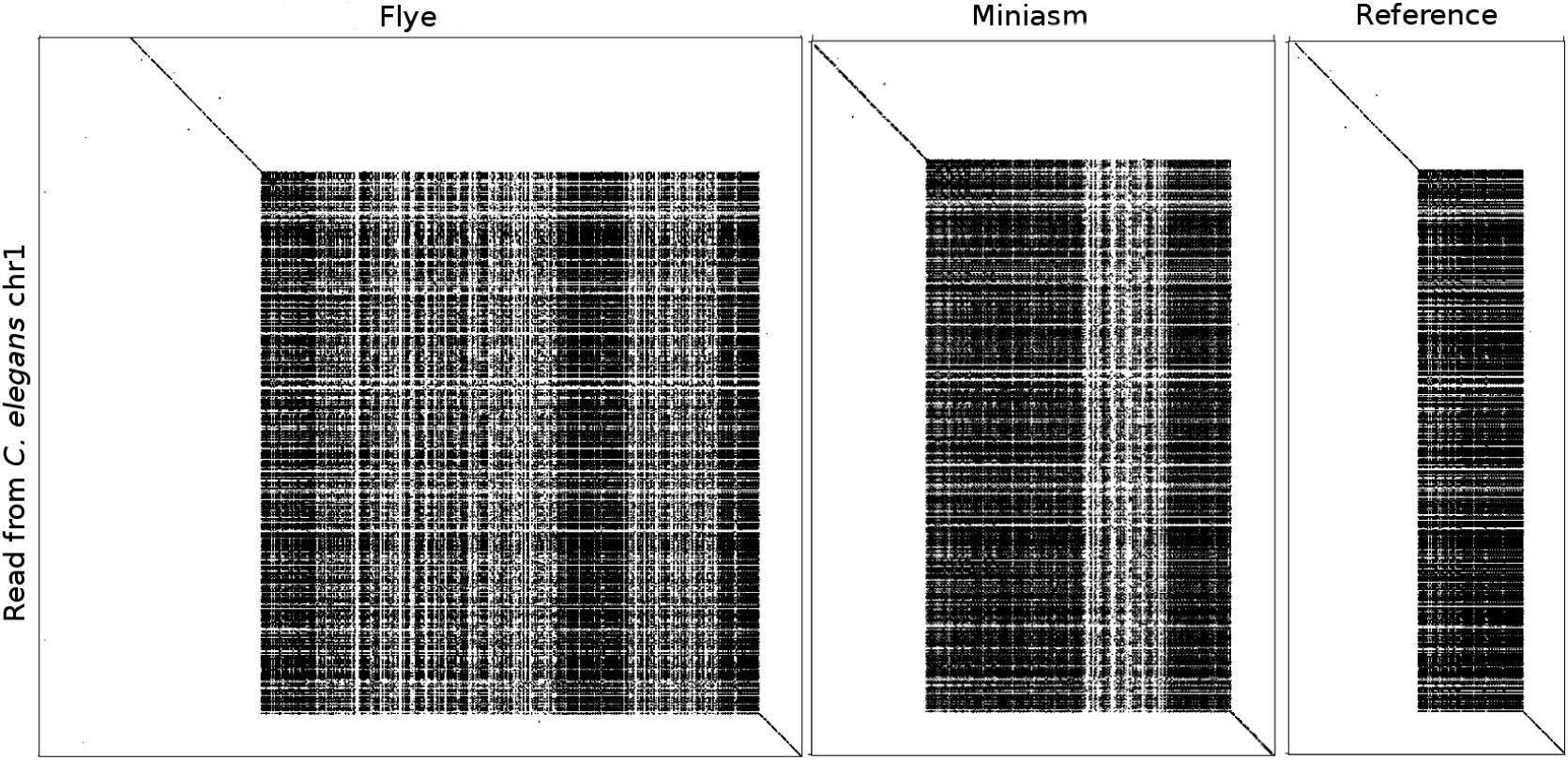
(Case B) Dot-plots showing the alignment of a read from chromosome 1 against the Flye assembly (left), the Miniasm assembly (middle) and the reference genome of *C. elegans* (right). The reference genome contains a tandem repeat cluster of length 2 kb on chromosome 1. In contrast, the Flye and Miniasm assemblies of this region suggest a tandem repeat of length 10 kb and 5.6 kb, respectively. A single read that spans over the tandem repeat supports the Flye assembly. Since the mean read length in the WORM dataset is 11 kb, it is expected to have a single read spanning a given 10.0 kb region, but many more reads spanning any 5.6 kb region (Miniasm assembly) or 2.0 kb region (reference genome). Six out of 23 reads cross the “left” border (two out of 18 reads cross the “right” border) of this tandem repeat by more than 5.6 kb, thus contradicting the length estimate given by Miniasm.

**Figure A8.**
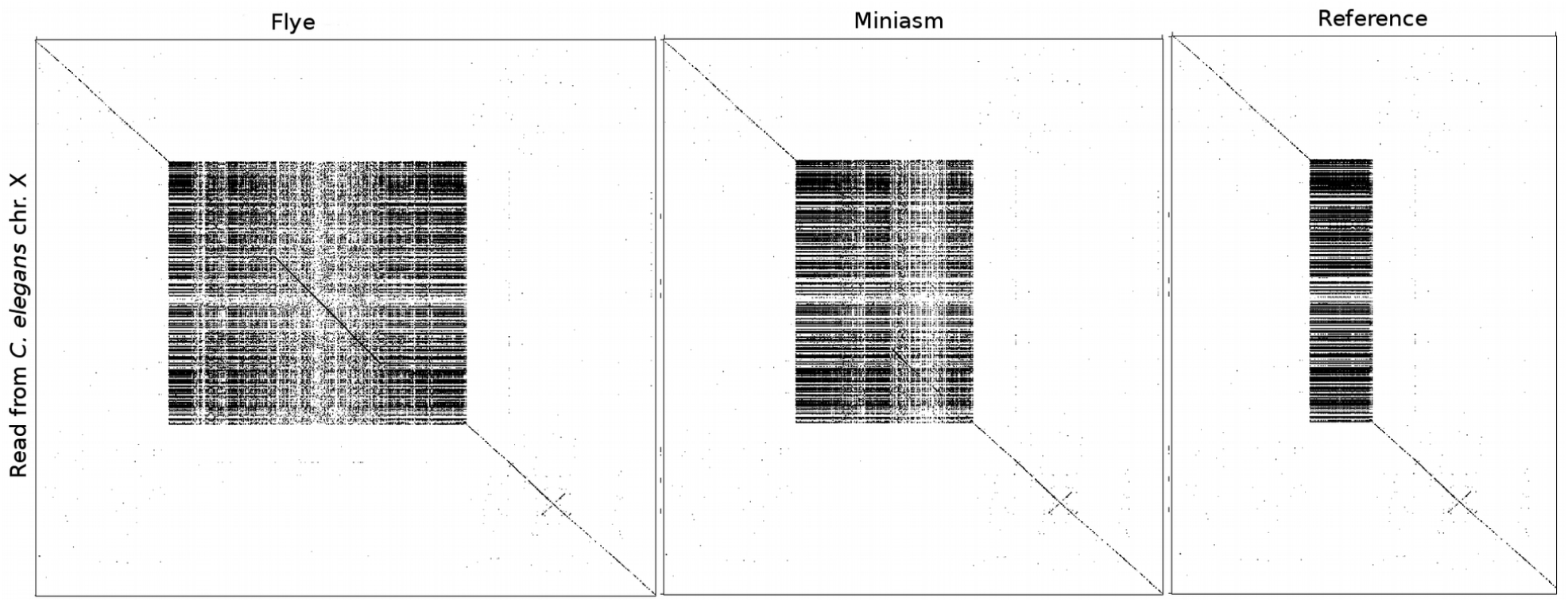
(Case C) Dot-plots showing the alignment of a read from chromosome X against the Flye assembly (left), the Miniasm assembly (middle) and the reference genome of *C. elegans* (right). The reference genome contains a tandem repeat of length 3 kb on chromosome X. In contrast, the Flye and Miniasm assemblies of this region suggest a tandem repeat of length 13.6 kb and 8 kb, respectively. A single read that spans over the tandem repeat reveals the repeat cluster to be of length 12.2k, which suggests that the Flye length estimate was more accurate.

**Figure A9.**
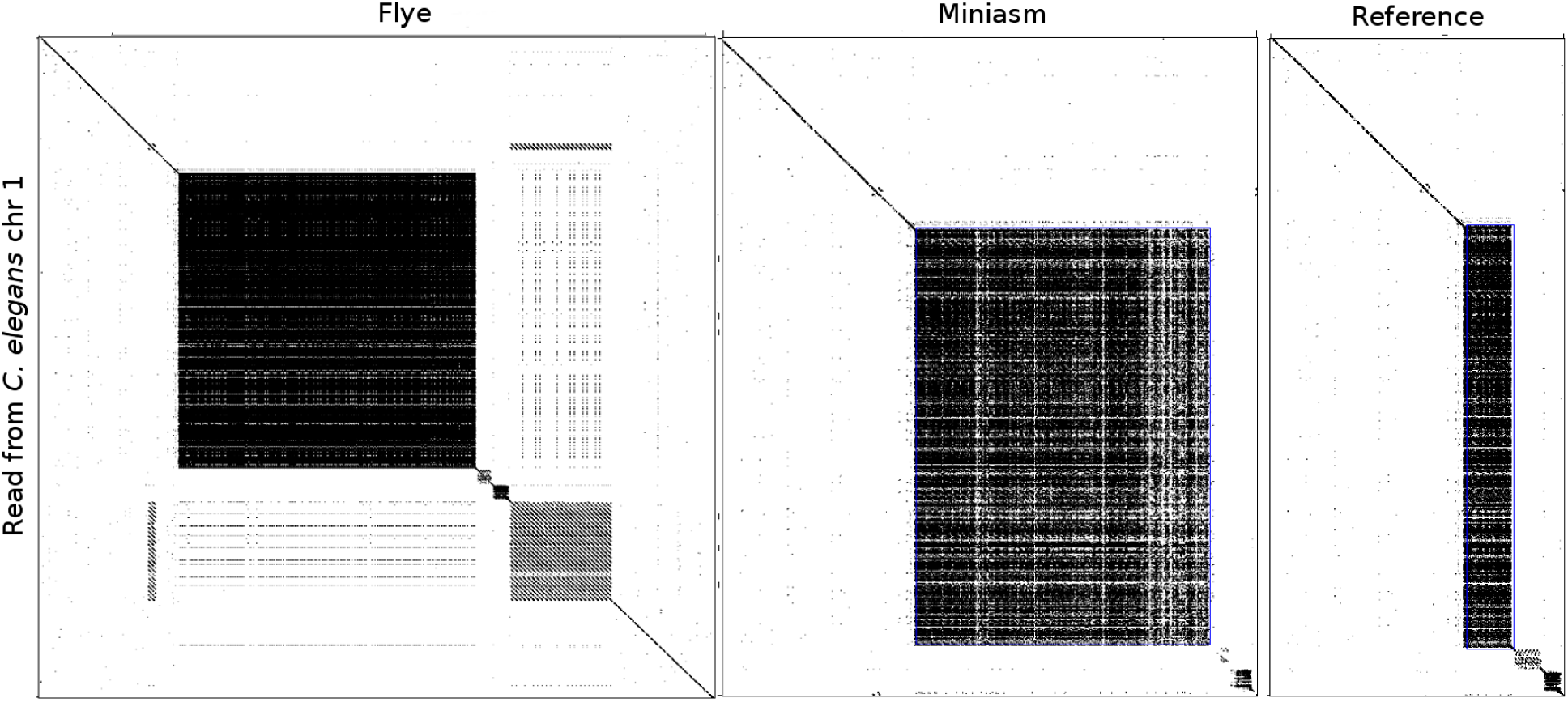
(Case D) Dot-plots showing the alignment of a read from chromosome 1 against the Flye assembly (left), the Miniasm assembly (middle) and the reference genome of *C. elegans* (right). The reference genome contains a tandem repeat of length 1.5 kb on the chromosome 1. In contrast, the Flye and Miniasm assemblies of this region suggest a tandem repeat of length 14 kb and 9.7 kb, respectively. Three reads that span over the tandem repeat reveal the repeat cluster to be of length 14k, which suggests that the Flye length estimate is more accurate.

**Figure A10.**
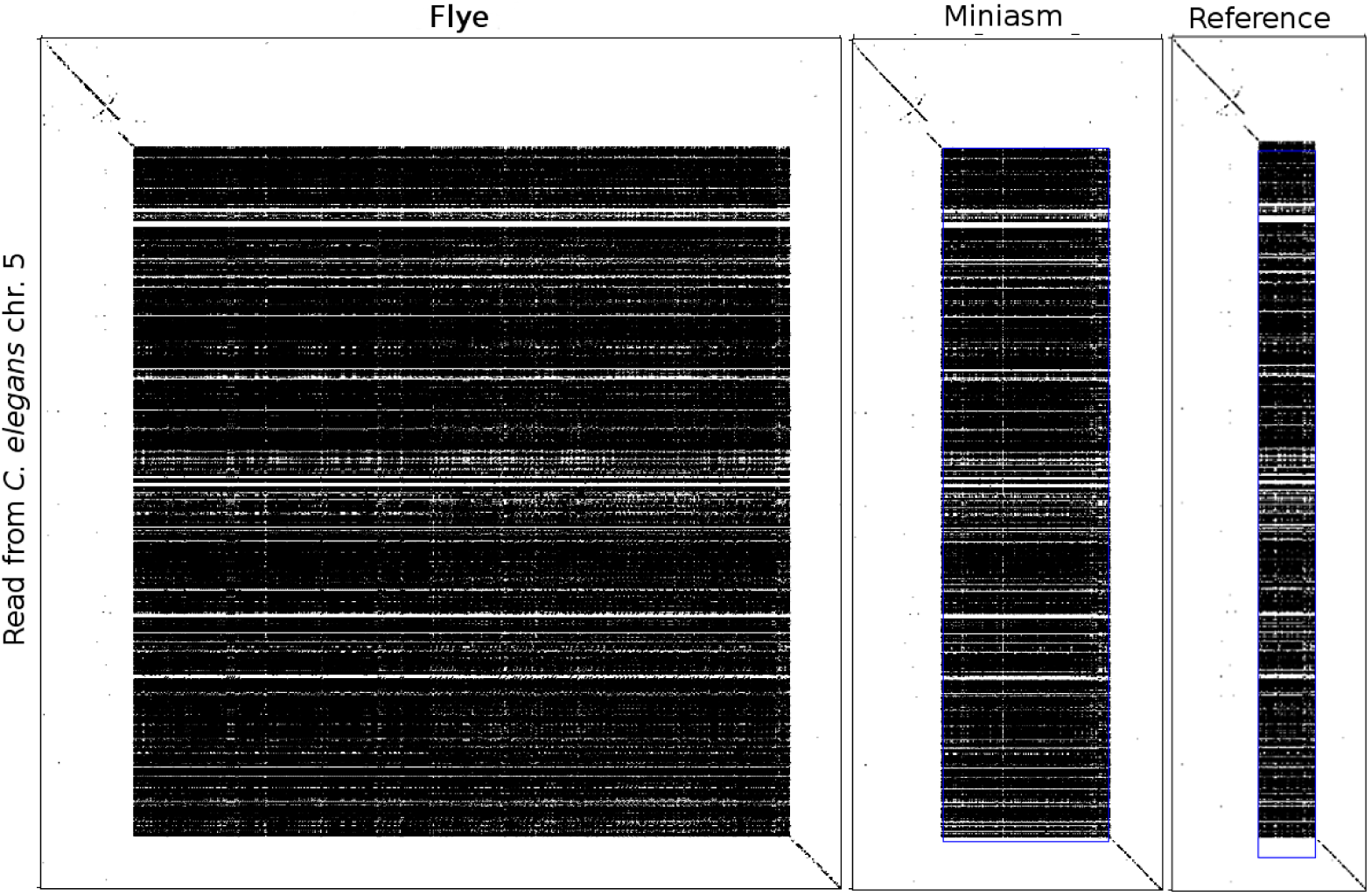
(Case E) Dot-plots showing the alignment of a read from chromosome 5 against the Flye assembly (left), the Miniasm assembly (middle) and the reference genome of *C. elegans* (right). The reference genome contains a tandem repeat of length 1.5 kb on the chromosome 1. In contrast, the Flye and Miniasm assemblies of this region suggest tandem repeat clusters of length 17 kb and 4.3 kb, respectively. One read that spans over the tandem repeat reveals the repeat cluster to be of length 18.0 kb, which suggests that the Flye length estimate was more accurate.

